# Clinical cure of chronic hepatitis B is dependent on activation and perpetuation of robust CD4^+^ T cell responses

**DOI:** 10.1101/2025.09.29.677401

**Authors:** Jillian M. Jespersen J, Lia Avanesyan, Jean Publicover, Nicholas D. Carey, Ravi K. Patel, Austin W. Edwards, Sarah Stenske, Jae Shin, Jiajing Li, Margaret Simone, Nayvin W. Chew, Nicholas Wong, Suprita Trilok, Ann Erickson, Arjun Rao, Christopher P. Loo, Michael Stec, Mark Anderson, Gavin Cloherty, Monika Sarkar, Alexis J. Combes, Adil E. Wakil, Mark R. Segal, Gabriela K. Fragiadakis, Stewart Cooper, Jody L. Baron

## Abstract

Chronic infection with hepatitis B virus (HBV), a major global pathogen, often leads to immune-mediated progressive liver injury and liver cancer. While seroclearance of the surface antigen (HBsAg) defines clinical cure and reduces disease-associated risks, HBsAg clearance is rarely observed and remains therapeutically elusive. Here we overcome some of the challenges to studying immune mechanisms of HBsAg clearance in chronic hepatitis B (CHB) using our mouse model of age-dependent HBsAg clearance and persistence, and samples from our BeNEG-DO clinical trial that provided longitudinal PBMCs from patients who either cleared HBsAg or retained stable HBsAg levels after stopping nucleos(t)ide analog therapy. We show that young mice fail to clear HBsAg and have impaired ability to efficiently initiate and sustain HBV-specific CD4^+^ T cell responses. We also demonstrate a role for CD4^+^ T cells in hepatic leukocyte organization and cytotoxicity, and in HBV-specific CD8^+^ T cell cytotoxicity and HBsAg clearance. Upstream of the CD4^+^ T cell response, we reveal that hepatic dendritic cells, particularly cDC2s, direct effective CD4^+^ T cell activation and differentiation. Studies in CHB patients identified immune features of HBsAg clearance that overlap with the mouse model, including T_H_1 and cytotoxic CD4^+^ T cell activation and CD8^+^ T cell cytotoxic effector function. These findings identify an essential role for potent CD4^+^ T cell activation in the clinical cure of CHB and illuminate potential immunotherapeutic targets for enhancing CD4^+^ T cell responses to achieve greater HBsAg clearance rates.

**One Sentence Summary:** Using a mouse model of hepatitis B and longitudinal PBMCs from patients with chronic hepatitis B, we identify shared mechanisms of HBsAg seroclearance.

## INTRODUCTION

Approximately 250 million people are chronically infected with hepatitis B virus (HBV) (*1*), a diagnosis established by persistence of the surface antigen (HBsAg) in serum. In 2022, chronic hepatitis B virus (CHB) was associated with approximately 1.1 million attributable deaths, a number predicted to increase through 2030 (*1, 2*). While most adults acutely infected with HBV launch an immune response that clears all HBV antigens and DNA from serum, most neonates and infants fail to clear HBsAg and remain persistently infected throughout life (*3, 4*) contributing an estimated 82% of the global disease burden of CHB (*5*). In CHB, ineffective immune responses directed at HBV-infected hepatocytes rarely clear HBsAg and often cause progressive liver injury leading to cirrhosis and higher risk for hepatocellular carcinoma (*6*). While the ineffective immunity of CHB can be dampened using nucleos(t)ide analogs (NAs) to suppress HBV DNA synthesis from pregenomic RNA (pgRNA), NAs seldom result in HBsAg seroclearance, the endpoint defining clinical (or “functional”) cure. Limitations of NAs mainly stem from persistence of transcriptionally capable covalently closed circular DNA and unrestrained HBsAg transcription from integrant HBV DNA (*7*). Therefore, standard-of-care NA therapy is usually indefinite, incurring cost, potential side effects, and patient concerns (*8, 9*). These aggregated factors establish urgent need to develop finite therapies for CHB that can achieve high rates of durable HBsAg clearance, for example by stimulating an appropriate immune response.

The inducibility of HBsAg clearance in CHB patients was initially established by adoptive transfer of donor immunity to bone marrow transplant recipients (*10, 11*), and more recently in some patients by activation of autologous immunity after NA withdrawal under specific protocols (*12*). To date, however, approaches aimed at overcoming the ineffective responses of CHB by manipulating autologous innate and adaptive immunity have been inefficient (*13*), highlighting an inadequate understanding of the elements required to induce an effective anti-HBV immune program. Deciphering these immunological programs and mechanisms of HBsAg clearance in CHB has been hampered by a paucity of experimental models and clinical samples for systematic study. Here, we overcome some of these challenges by using our mouse model of predictable HBsAg clearance and persistence (*14, 15*) along with patient samples from our BeNEG-DO clinical trial (NCT02845401). The mouse model provided a vehicle to identify components of hepatic immunity that influence HBsAg outcome, and our clinical trial provided longitudinal PBMCs from CHB patients who either cleared HBsAg or retained stable HBsAg levels.

The mouse model mirrors important immunological aspects of human HBV infection including the impact of host age on HBsAg outcome (*15, 16*). In this model, HBV transgenic C57BL/6 mice with hepatic expression of either select HBV antigens or infectious genotype D human HBV are crossed to a RAG1-deficient background (*Rag1*^-/-^) to prevent the development of an adaptive immune system that would otherwise be tolerant to HBV antigens (HBVtgRag^−/−^ mice). Adoptive transfer of HBV-naïve splenocytes from adult wild-type mice reconstitutes the recipient adaptive immune system and initiates an anti-HBV immune response. After reconstitution, adult HBVtgRag^−/−^ recipient mice generate a diverse HBV-specific T cell response and clear circulating HBsAg which is accompanied by a serological profile of HBV core antibody (HBcAb)^+^, HBsAg^−^, anti-HBs^+^, phenocopying patients who clear acute HBV infection (*15*). Conversely, transfer of splenocytes into young HBVtgRag^−/−^ mice leads to a weaker acute response and the serological profile of persons who develop CHB (HBcAb^+^, HBsAg^+^, anti-HBs^−^). This model has allowed interrogation of hepatic immune mechanisms associated with HBsAg clearance or persistence and the development of hypotheses to be tested in patients (*15–17*).

Complementing our studies in this model, the BeNEG-DO clinical trial provided our first gateway to study CHB patients who either cleared HBsAg or retained stable HBsAg levels after stopping NAs. The trial was initiated after a landmark study in e-Antigen negative adult CHB patients demonstrated that unleashed HBV replication after withdrawing ≥192 weeks of NA therapy could induce HBsAg clearance in >25% of patients (*12*) – establishing that effective immunity can be stimulated in adults despite their history of futile immunity, often since the neonatal period. Our clinical trial (NCT02845401) is a prospective case-control trial of similar design, involving 76 predominantly Asian CHB cases in the San Francisco Bay area who stopped NA after at least 192 weeks of continuous therapy. The trial was designed not only to verify safety and benefit of timed treatment withdrawal in our population but also to seek immunological features associated with HBsAg outcomes using PBMCs obtained before, and at clinically distinct timepoints after NAs were stopped.

These studies identified shared features of HBsAg clearance in the mouse model and human patients, including a pivotal role for T_H_1 and cytotoxic CD4^+^ T cell activation, and CD8^+^ T cell cytotoxic effector function. The mouse studies illuminated the hepatic immune orchestration of HBsAg clearance, which involved differences in the antigen presentation capacity of dendritic cells and hepatic leukocyte organization. The mouse and human data also suggest that generation and perpetuation of an adaptive HBV immune response with minimal capacity for CD4^+^ T cell activation facilitates HBsAg persistence. Our studies newly identify immune programs associated with effective and ineffective immunity in CHB that can have translational significance.

## RESULTS

### Hepatic HBsAg-specific CD4^+^ T cell activation is impaired in young mice that fail to clear HBsAg

Because our prior work pointed to a pivotal role for CD4^+^ T follicular helper (T_FH_) cell activation in HBsAg clearance (*15*), we investigated hepatic CD4^+^ T cell activation, differentiation, and functional capacity in young versus adult recipient mice. Using spectral flow cytometry, including two HBsAg-specific MHCII tetramers, we studied total and HBsAg-specific intrahepatic CD4^+^ T cells in adult and young mice at clinically relevant phases after adoptive transfer: initial priming (days 3-5), peak immune activation (day 8), contraction and resolution of hepatitis (day 15), HBsAg clearance/persistence (day 30), and memory/chronicity (day 60+) (Fig. 1A, fig. S1A-C).

**Fig. 1:**
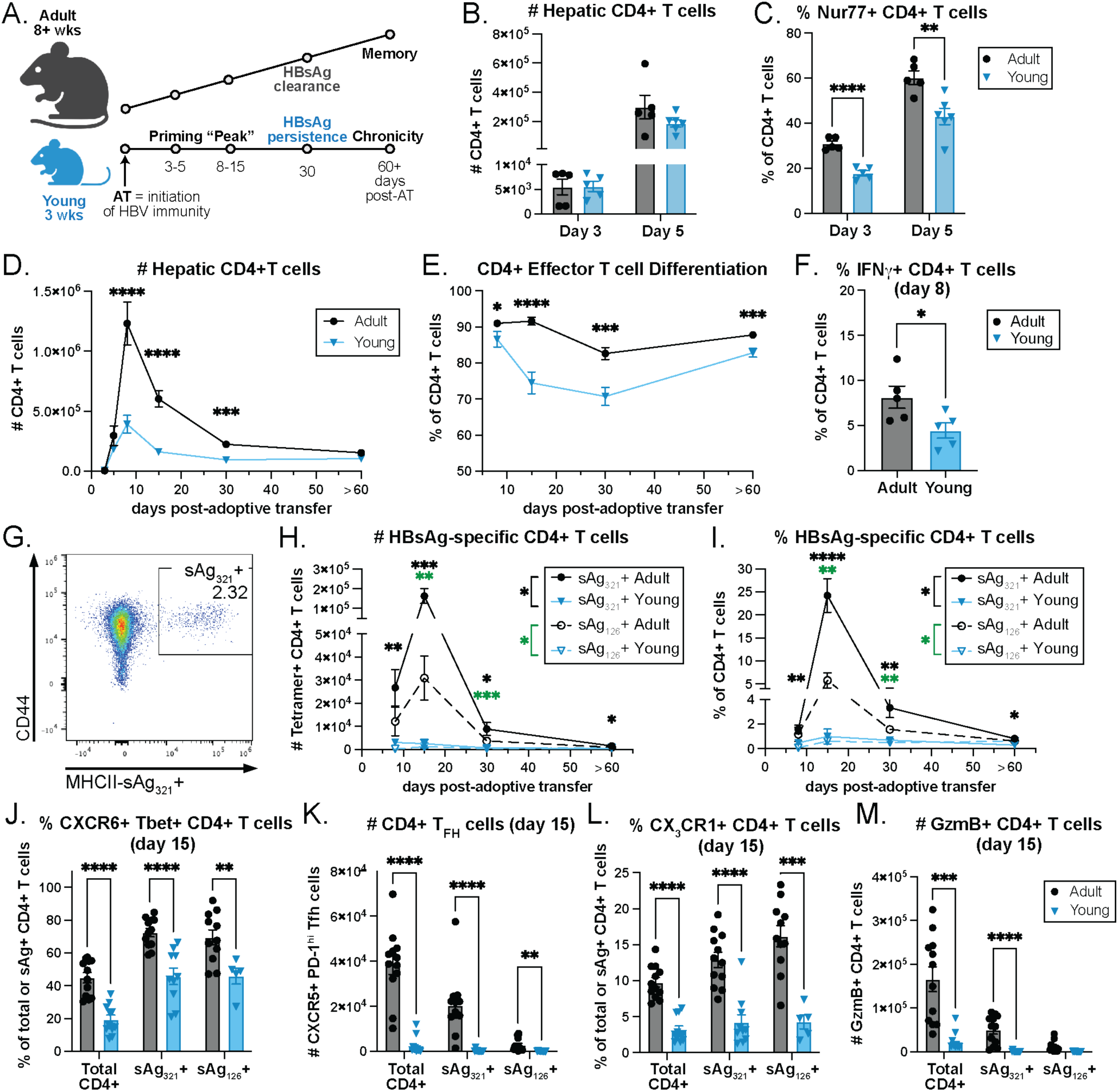
Hepatic CD4+ T cell activation and differentiation, but not initial recruitment, is dependent on the age of the liver microenvironment. (**A**) Adult (> 8 wks) and young (3-3.5 wks, pre-weaning) HBVtgRag-/- mice adoptively transferred on day “0” with WT splenocytes and hepatic leukocytes were assessed by flow cytometry at the indicated time points. (**B**) Total number of hepatic CD4+ T cells on day 3 and 5 post-adoptive transfer in adult (black circles) and young (blue triangles) mice. (**C**) Percentage of CD4+ T cells that are Nur77+. (**D**) Total number of hepatic CD4+ T cells on days 3, 5, 8, 15, 30, and ≥60 post-adoptive transfer. (**E**) Percentage of total CD4+ T cells with an effector/effector memory phenotype; CD44+ CD62L-. (**F**) Percentage of hepatic CD4+ T cells that are IFNγ+ on day 8. (**G-I**) HBsAg-specific CD4+ T cells were identified with MHCII-sAg_321-335_ and MHCII-sAg_126-138_ tetramers. (**G**) Representative flow plot demonstrating sAg_321_ tetramer staining. (**H**) Total number and (**I**) percentage of hepatic sAg_321_-specific (solid lines) and sAg_126_-specific (dashed lines) CD4+ T cells. Percentage of total or HBsAg-specific hepatic CD4+ T cells that are (**J**) CXCR6+ Tbet+ or (**L**) CX_3_CR1+. Number of (**K**) CXCR5+ PD-1+ CD4+ T_FH_ cells and (**M**) GzmB+ CD4+ T cells. (B-D, F) N≥5 per group, per time point, data representative of at least two independent experiments or (H-M) N≥11 per group, per time point, combined from two-three independent experiments; bars represent mean ± SEM; statistics performed using unpaired t-test without multiple test correction when comparing two groups or Dunnett’s multiple comparison test when comparing more than two groups; * p<0.05, ** p<0.01, *** p<0.001, **** p<0.0001; black * indicate comparisons between sAg_321_+ and green * indicate comparisons between sAg_126_+ populations.

After adoptive transfer into young or adult HBVtgRag^−/−^ mice, initial CD4^+^ T cell infiltration into the liver was equivalent, independent of age. Few CD4^+^ T cells were detectable on day 3 whereas CD4^+^ T cells expanded >40-fold in both adult and young livers by day 5 (Fig. 1B, fig. S1D). Although early hepatic infiltration was not different by age, CD4^+^ T cells in adult livers showed signs of increased TCR activation demonstrated by higher expression of Nur77, an immediate-early gene rapidly upregulated upon TCR engagement (Fig. 1C) (*18*). Hepatic CD4^+^ T cell expansion continued through day 8 and began to contract by day 15. The magnitude of CD4^+^ T cell expansion within the liver was substantially greater in adult mice compared to young mice and persisted through the antigen clearance phase (day 30) before leveling out to similar numbers of total CD4^+^ T cells by day 60 (Fig. 1D, fig. S1D). These data suggest age-dependent differences in the capacity of the liver microenvironment to prime and support CD4^+^ T cells. In addition to the more robust expansion and activation of hepatic CD4^+^ T cells in adult recipient mice during the acute phase response, these CD4^+^ cells exhibited increased effector T cell differentiation (CD44^+^ CD62L^-^) and secretion of IFNγ (Fig. 1E and F).

To examine HBV antigen specificity among liver infiltrating CD4^+^ T cells, we generated two I-A(b) MHCII tetramers folded with HBsAg peptide epitopes: sAg_321-335_ (FGKFLWEWASARFSW), identified by ELISpot analysis in our laboratory, and a known epitope, sAg_126-138_ (RGLYFPAGGSSSG) (*19*). Beginning on day 8, tetramer-stained CD4^+^ T cells became detectable and were substantially enriched in adult compared to young mice throughout the response into the memory phase (Fig. 1G-I, fig. S1E). In young mice, hepatic tetramer-positive events were at the threshold of detection on day 8, became detected at low frequency on day 15, and at all subsequent timepoints returned to near the threshold of detection. Thus, while young mice may have a limited capacity to prime HBsAg-specific CD4^+^ T cells, the rare nature of these cells suggests that the young liver microenvironment fails to efficiently promote and sustain an effective anti-HBsAg CD4^+^ T cell response.

Antigen-specific CD4^+^ T cells also demonstrated significant age-dependent phenotypic differences including T_H_1 differentiation, effector cytokine production, and tissue residency (Fig. 1J-M, fig. S2). Specifically, a greater amount of total and HBsAg-specific hepatic CD4^+^ T cells from adult livers were Tbet^+^ CXCR6^+^ T_H_1 effector cells compared to CD4^+^ T cells from young livers (Fig. 1J, fig. S1F and G, fig. S2B). The percentage of HBsAg-specific CD4^+^ T cells that had differentiated into T_FH_ cells was not different by age suggesting that T_FH_ differentiation can occur in the young liver (fig. S1H). However, the total number of hepatic HBsAg-specific T_FH_ cells was substantially increased in adult livers, consistent with our previous findings that T_FH_ cells are expanded in adult recipient mice and crucial for initiating HBsAg clearance and anti-HBs production (Fig. 1K) (*15*). Notably, the expression of CX_3_CR1, a marker of terminal differentiation enriched in cytotoxic T lymphocytes (CTL) (*20*), together with Granzyme B and Perforin, demonstrates that effective CD4^+^ T cell priming in the adult mice also includes the expansion of cytotoxic CD4^+^ T cells (Fig. 1L and M, fig. S1I-L, fig. S2C-D). Together these results reveal that compared with adult mice, hepatic CD4^+^ T cell priming, differentiation, and functional capacity is markedly impaired in young livers, including a stunted HBsAg-specific CD4^+^ effector and memory T cell response.

### HBsAg clearance is associated with CD4^+^ T cell-rich hepatic leukocyte clusters

To assess hepatic positioning, organization, and in situ interactions of T cells during peak inflammation, we performed immunohistochemistry to identify CD4^+^ and CD8^+^ T cells in the livers of adult and young mice on day 8 (Fig. 2). In adult livers, leukocyte clusters (>5 cells within 100 pixels, see supplemental materials and methods) were larger in size and consisted of both CD4^+^ and CD8^+^ lymphocytes, compared to smaller clusters that principally consisted of CD8^+^ lymphocytes in young livers (Fig. 2A and B). However, while we observed an age-dependent increase in the number of hepatic CD4^+^ T cells in adult mice, we did not detect a difference in the number of CD8^+^ T cells (Fig. 2C). Furthermore, a greater percentage of the hepatic lymphocytes were associated with an immune cluster in adult compared with young mice and the size of those clusters were larger (Fig. 2D-E). Hepatic immune clusters in adult mice that clear HBsAg on average had 1.6x more cells per cluster compared to young mice (Fig. 2E).

**Fig. 2:**
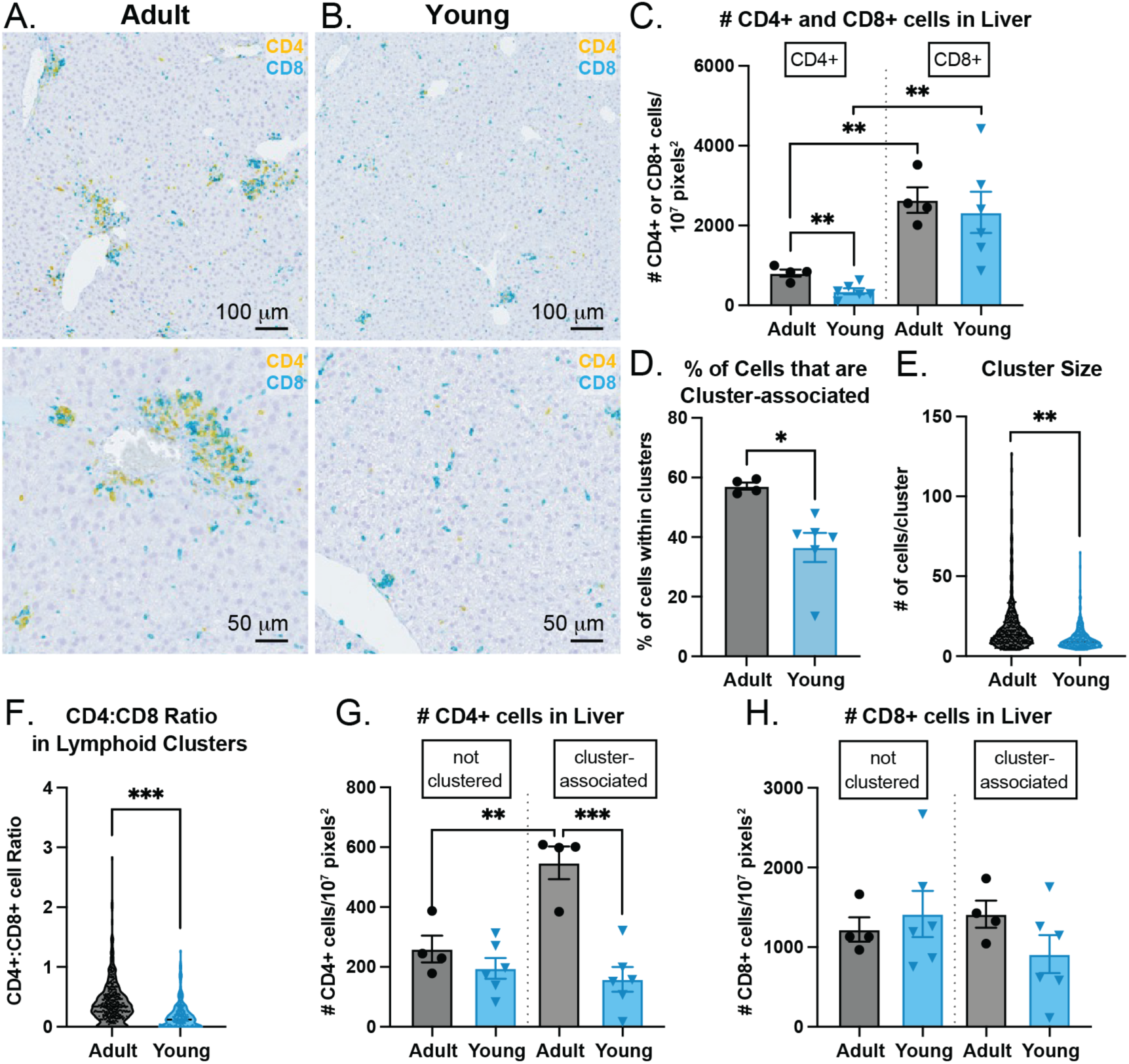
CD4+ lymphocytes are enriched in hepatic leukocyte clusters associated with HBsAg clearance. (**A-B**) Representative images of FFPE liver tissue from (**A**) adult and (**B**) young HBVtgRag-/- mice on day 8 stained with anti-CD4 (yellow), -CD8 (teal), and RORγ/γ(t) (purple). (**C**) Number of CD4+ and CD8+ cells in adult (black) and young (blue) livers normalized to tissue area (pixels^2^). (**D**) Percentage of cells associated with a cluster (defined as ≥5 CD4+, CD8+, and/or RORγ+ cells within 100 pixels or ∼22 mm). (**E**) Number of cells present within each cluster. (**F**) Ratio of CD4+ to CD8+ cells within leukocyte clusters. (**G**) Number of hepatic CD4+ or (**H**) CD8+ cells located within a cluster or individually (“not clustered”). Bars represent mean ± SEM; statistics performed using (C-D, G-H) unpaired t-test or (E-F) linear mixed model to account for sample-level variation.

Compositionally, although CD8^+^ cells dominated hepatic immune clusters of both young and adult mice, the ratio of CD4^+^:CD8^+^ cells shows significant enrichment of CD4^+^ cells in the clusters from adult compared to young mice (Fig. 2F). Further, this enrichment of hepatic CD4^+^ cells specifically occurred within immune clusters, suggesting local expansion or recruitment, and was not observed for CD8^+^ cells (Fig. 2G and H). Taken together, these data support prior work by us and others highlighting a conditional role for the liver as a site of productive antigen presentation and T cell priming (*17, 21–25*), and further suggest that these hepatic immune clusters serve as an essential priming hub or immune scaffold that can facilitate efficient immune activation, expansion, and cross-talk necessary for the effective CD4^+^ and CD8^+^ T cell responses required for HBsAg clearance.

### Hepatic dendritic cells drive effector differentiation and function of CD4^+^ T cells

To understand the upstream events that influence hepatic T cell responses and ultimately HBsAg clearance and seroconversion, we analyzed the professional antigen presenting cells (APCs) in the liver – Kupffer cells (KCs), monocyte-derived macrophages (MDM≥), and dendritic cells (DCs) – to identify age-dependent differences in the immune microenvironment upon initiation of the adaptive immune response. At baseline, before adoptive transfer, adult HBVtgRag-/- mice, compared to young mice, had a greater abundance of type 2 conventional DC (cDC2s), cells known for their capacity to prime CD4^+^ T cell responses (Fig. 3A, fig. S3C) (*26*). There were no significant age-dependent differences in the abundance of cDC1s, KCs, or MDM≥ (Fig. 3A, fig. S3A and B). The hepatic cDC2 population expanded dramatically in adult mice by day 8 and remained differentially enriched in an age-dependent manner into the late memory phase when “young” mice have matured into adulthood, suggesting durable changes to the DC population following their futile immune response (Fig. 3A). Notably, experiments using CD45.1^+^ congenic splenocytes adoptively transferred into CD45.2^+^ HBVtgRag^−/−^ recipient mice demonstrated that this increase in hepatic cDC2s was entirely attributed to expansion and/or differentiation of host-derived (CD45.2^+^) cells, rather than infiltration or differentiation of transferred donor splenocytes (fig. S3D). The age-dependent difference in cDC2 abundance was also observed in WT C57BL/6 mice, suggesting this difference is physiological, rather than a property restricted to HBVtgRag^−/−^mice (fig. S3C and F).

**Fig. 3:**
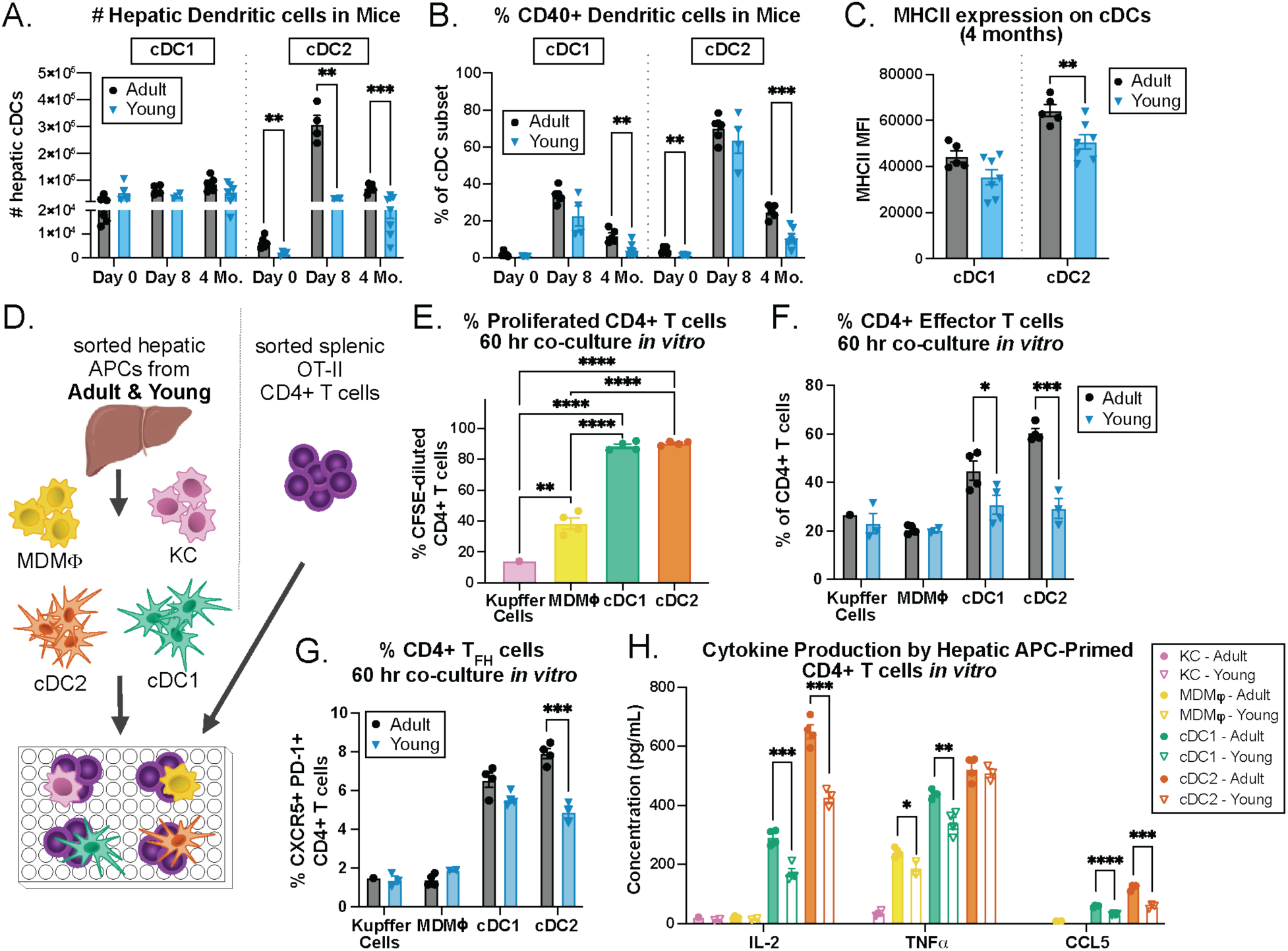
Age-dependent antigen presentation by hepatic dendritic cells drives effector differentiation and function of CD4+ T cells. (**A**) Number of hepatic adult (black circles) or young (blue triangles) cDC1 (NK1.1- Ly6G- F480- CD11b- MHCII+ CD11c+ XCR1+ CD172a-) and cDC2 (NK1.1- Ly6G- F480- MHCII+ CD11c+ XCR1-CD172a+). (**B**) Percentage of CD40+ cDC1 and cDC2. (**C**) Median fluorescence intensity (MFI) of MHCII on cDC1 and cDC2. (**D**) Hepatic monocyte-derived macrophages (MDM≥; CD45+ NK1.1- CD90.2- MHCII+ CD11c - F4/80+ CD64+ CD11b^hi^ TIM4^lo^); Kupffer cells (KC; CD45+ NK1.1- CD90.2- MHCII+ CD11c- F4/80+ CD64+ CD11b^lo^ TIM4^hi^); cDC1 (CD45+ NK1.1- CD90.2- F4/80- CD64- MHCII+ CD11c+ XCR1+ CD172a-); and cDC2; (CD45+ NK1.1- CD90.2- F4/80- CD64-MHCII+ CD11c+ XCR1- CD172a+) were FACS-sorted, pulsed with OVA_323-339_ peptide, and cocultured with CFSE-labeled splenic CD4+ OT-II cells for 60 hrs. (**E**) Percentage of CD4+ T cells co-cultured with adult hepatic myeloid cells that underwent proliferation (CFSE diluted). Percentage of CD4+ T cells with an (**F**) effector (CD44+ CD62L-) and (**G**) T follicular helper cell (T_FH_; CXCR5+ PD1+) phenotype. (**H**) Concentration of IL-2, TNFα, and CCL5 in coculture supernatants. (A-C) N≥4, representative of 2 or more independent experiments, (E-F) N=pooled from ≥8 mice per group, dots represent technical replicates; bars represent mean ± SEM; statistics performed using unpaired t-test without multiple test correction.

In addition to age-dependent differences in hepatic cDC2 abundance, we found evidence of increased intrinsic antigen presentation capacity of both hepatic cDC2s and cDC1s from adult mice compared to young mice, as demonstrated by increased expression of MHCII and the co-stimulatory molecules CD40 and CD86 (Fig. 3B and C, fig. S3E and G). To determine whether hepatic DCs have a differential capacity by age to activate T cells, we developed an *in vitro* CD4^+^ T cell stimulation assay using hepatic APCs. CD4^+^ OT-II splenocytes from TCR transgenic mice were co-cultured with equivalent numbers of either adult or young sorted hepatic APCs loaded with cognate antigen (ovalbumin peptide, Ova_323-339_), followed by assessment of T cell activation and cytokine secretion (Fig. 3D). As expected, these experiments showed a heightened ability of DCs over other hepatic APCs to stimulate CD4^+^ OT-II cell proliferation, with 90% of naïve CD4^+^ OT-II cells proliferating when primed with either cDC1s or cDC2s, while only 40% and 15% of cells proliferated when primed by MDM≥ and KCs, respectively (Fig. 3E). Furthermore, we found no age-dependent difference in the ability of hepatic MDM≥ or KCs to induce differential activation or differentiation of CD4^+^ T cells *in vitro* (Fig. 3F and G).

In contrast, hepatic cDC2s from adult mice, and to a lesser degree cDC1s, were stronger drivers of T_EFF_ and T_FH_ differentiation compared with DCs from young mice, similar to our observations of age-dependent CD4^+^ T cell responses against HBV *in vivo* (Fig. 3F and G). This age-dependent difference in DC capacity to activate CD4^+^ T cells was specific to liver DCs, since splenic DCs from both adult and young mice led to equivalent CD4^+^ T_EFF_ and T_FH_ differentiation (fig. S4A-C). Hepatic DCs also showed differential capacity by age to induce cytokine and chemokine production within these co-cultures, with notable increases in IL-2, TNFα, and CCL5 secretion when CD4^+^ T cells were activated by adult cDCs, especially cDC2s, which showed the strongest activation potential of all tested populations (Fig. 3H). Notably, KCs and hepatic MDM≥ had little capacity to induce these secreted factors. Taken together, these data suggest a model in which hepatic adult cDC2s, which are both more abundant and intrinsically superior APCs within the liver microenvironment compared to their young mouse counterparts, support efficient priming, activation, and differentiation of naïve CD4^+^ T cells, which in turn supports hepatic immune aggregates and immune cell crosstalk, ultimately resulting in HBsAg clearance and anti-HBs production.

### CD8^+^ T cell responses are transiently impaired in young mice that fail to clear HBV

To complement studies of HBsAg-specific CD4^+^ T cell responses in our model, we identified two MHC class I (H2-Kb)-restricted HBsAg epitopes to study the synchronous CD8^+^ T cell responses: sAg_353-360_ (VWLSVIWM) and sAg_370-378_ (SILSPFLPL) (Fig. 4A) (*27*). Similar to the HBsAg-specific CD4^+^ T cells, hepatic tetramer^+^ CD8^+^ T cells became detectable at day 8 post-adoptive transfer (Fig. 4B, fig. S5A). However, unlike HBsAg-specific CD4^+^ T cells in young mice, tetramer^+^ CD8^+^ T cell populations were readily detected in young livers by day 15 and continued to expand over time. In adult mouse livers between day 8 and day 30, when HBsAg clearance is occurring, there was a significantly higher frequency of antigen-specific CD8^+^ T cells compared to young mice; the response demonstrating epitope-dependent expansion characteristics. While the sAg_353_ response peaks early then plateaus, the sAg_370_ response was relatively delayed and continued to expand over time, both in absolute numbers and as a proportion of the total CD8^+^ T cell compartment, with no apparent contraction phase (Fig. 4B, fig. S5A). At all later timepoints analyzed, including the memory phase beyond 60 days, the HBV-specific CD8^+^ T cell response in “young” mice was quantitatively robust and largely equivalent to adult counterparts. This HBV-specific CD8^+^ T cell response in young mouse livers contrasts with our findings in the CD4^+^ T cell compartment, which produced no evidence in young recipient mice of a robust antigen-specific response or of an HBV-specific memory CD4^+^ T cell population (Fig. 1H and I).

**Fig. 4:**
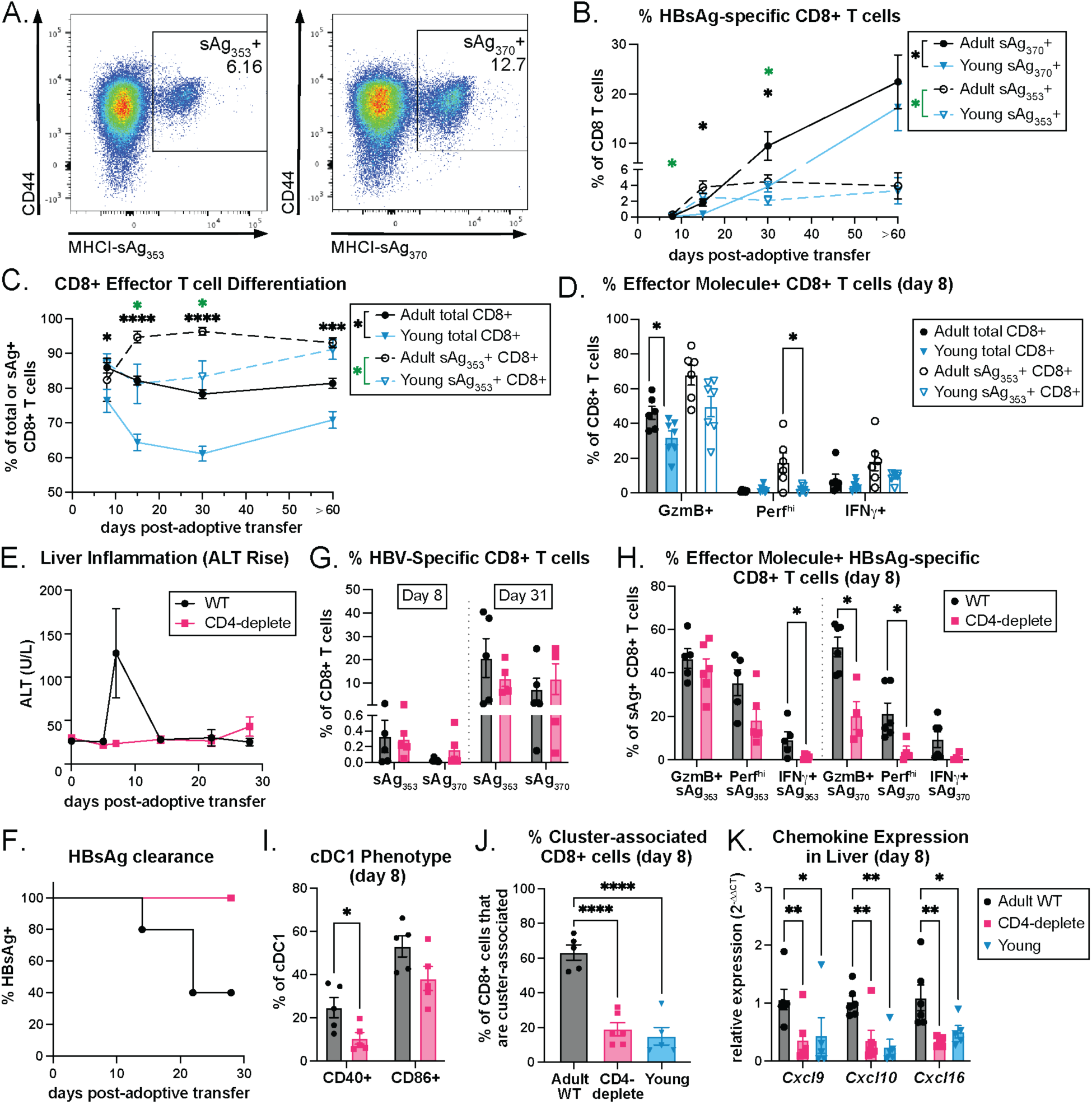
HBsAg clearance and effective CD8+ T cell function is CD4+ T cell-dependent. (**A**) Representative flow plots demonstrating staining with MHCI-sAg_353-360_ and MHCI-sAg_370-378_ tetramers. (**B**) Percentage of hepatic sAg_370_-specific (solid lines) and sAg_353_-specific (dashed lines) CD8+ T cells on days 8, 15, 30 and ≥60. (**C**) Percentage of total (solid) or sAg_353_-specific (dashed) CD8+ T cells with an effector/effector memory phenotype; CD44+ CD62L-. (**D**) Percentage of total (filled bars) or sAg_353_-specific (empty bars) CD8+ T cells expressing Granzyme B (GzmB), Perforin (Perf^hi^), and IFNγ on day 8. Adult mice that received WT splenocytes (WT; black circles) or WT splenocytes depleted of CD4+ cells (CD4-deplete; pink squares) were assayed for plasma (**E**) ALT and (**F**) HBsAg levels. (**G**) Percentage of sAg_353_+ and sAg_370_+ CD8+ T cells days 8 and 31. (**H**) Percentage of sAg_353_+ (left) and sAg_370_+ (right) CD8+ T cells that were GzmB+, Perf^hi^, and IFNγ+ on day 8. (**I**) Percentage of CD40+ and CD86+ cDC1. (**J**) Percentage of CD8+ cells that were associated with an immune cluster (≥4 CD8+ cells) by tissue staining of adult WT (black), adult CD4-deplete (pink), or young (blue) livers. (**K**) Expression of hepatic *Cxcl9, Cxcl10,* and *Cxcl16* mRNA transcripts relative to *Gapdh* measured by qPCR. (B-C) N≥9 per group, per timepoint, combined from two-three independent experiments, (D) N≥6, representative of two independent experiments; bars represent mean ± SEM; statistics performed using unpaired t-test without multiple test correction. (E-K) Day 8: N≥5, day 31: n≥4, representative of two independent experiments; bars represent mean ± SEM; statistics performed using (B-E, G-K) unpaired t-test without multiple test correction when comparing two groups or Dunnett’s multiple comparison test when comparing more than two groups, and (F) log-rank test.

Given these CD8^+^ T cell findings, we sought qualitative differences between the adult and young CD8^+^ T cells during the HBsAg clearance window. Hepatic HBsAg-specific CD8^+^ T cells in adult mice demonstrated increased effector differentiation on days 15 and 30 but were equivalent to young mice by the memory phase (Fig. 4C, fig. S5B). Furthermore, adult CD8^+^ T cells expressed higher levels of Granzyme B, Perforin, and IFNγ during the peak of the response on day 8 (Fig. 4D, fig. S5C, Fig. S6). Thus, while young mice have the capacity to mount a substantial HBsAg-specific CD8^+^ T cell response over time, a transient defect in their functional differentiation during a critical early phase of the response may play a decisive role in their failure to clear HBsAg. Of note, the HBsAg-specific CD8^+^ T cells did not exhibit age-dependent differences in classical signatures of exhaustion during the memory/chronicity phase, nor were there differences in expression of memory markers (TCF-1, CD127) or effector molecules (GzmB, Perf) at this time (fig. S5D-F).

### CD4^+^ T cells are required for effective CD8^+^ T cell function and positioning

Early studies of acute HBV in adult chimpanzees demonstrated that depletion of either CD8^+^ or CD4^+^ T cells during the initial phase of infection prevented HBsAg clearance, establishing that both T cell subsets contribute to effective HBV immunity (*28, 29*). The critical link between CD4^+^ T cell help and effective CD8^+^ T cell responses (*30*), and our finding that young mice have an impaired HBsAg-specific CD4^+^ T cell response (Fig. 1), motivated us to investigate the role of CD4^+^ T cell help in the resulting CD8^+^ T cell response and to changes in the liver immune environment including DC licensing. For these studies, we compared adult HBVtgRag^−/−^ mice adoptively transferred with either complete splenocytes or splenocytes depleted of CD4^+^ T cells (Fig. 4E-K). Like young mice, adult mice receiving CD4-depleted splenocytes did not exhibit hepatocyte injury (no ALT rise) and were unable to clear HBsAg, highlighting the essential role of CD4^+^ T cells in HBsAg clearance, as observed in chimpanzees (Fig. 4E and F, fig. S1A).

Furthermore, as shown in young mice, CD4^+^ T cell depletion did not prevent expansion of HBsAg-specific CD8^+^ T cells in adult mice, since there was no difference in the number of sAg_353_^+^ or sAg_370_^+^ CD8^+^ T cells on day 8 or 31 (Fig. 4G, fig. S4G), indicating that CD8^+^ T cell priming can occur independently of CD4^+^ T cell help. When deprived of CD4^+^ T cell help, however, these effector HBsAg-specific CD8^+^ T cells produced lower amounts of Granzyme B, Perforin, and IFNγ, indicating decreased cytotoxic effector function, reminiscent of the CD8^+^ T cell response in young mouse livers (Fig. 4H, fig. S4H).

Because activated CD4^+^ T cells contribute to DC licensing and generation of CD8^+^ CTL immunity, we studied markers of hepatic DC licensing in the CD4^+^ T cell depletion model. Although both groups of recipient adult mice initially have equivalent DC populations, the group depleted of CD4^+^ T cell help exhibited significantly lower surface expression of CD40 by hepatic cDC1s at day 8 after adoptive transfer, suggesting that these unlicensed APCs less efficiently cross-present antigen resulting in an impaired early CD8^+^ T cell effector response (Fig. 4I) (*31–33*). Notably, mice lacking CD4^+^ T cells also had reduced frequency of cDC2s with diminished expression of MHCII compared to mice with CD4^+^ T cells (fig. S4I-K), similar to the age-dependent differences observed in the livers of young versus adult mice (Fig. 3A and C). Depletion of CD4^+^ T cells also led to an altered distribution of CD8^+^ T cells in the liver. Like in young mice, we observed a significant decrease in the percentage of cluster-associated CD8^+^ T cells in the adult mice that lacked CD4^+^ T cells (Fig 4J, fig. S4L-N). Consistent with this finding, there was lower abundance of *Cxcl9*, *Cxcl10, and Cxcl16* chemokine transcripts in the livers of adult mice lacking CD4^+^ T cell help, similar to young mice that have CD4^+^ T cells (Fig. 4K). These experiments indicate that efficient CD4^+^ T cell help contributes to a local hepatic microenvironment capable of recruiting CXCR3^+^ and CXCR6^+^ lymphocytes to sites enriched in the costimulatory signals necessary for effective activation and functional differentiation that can lead to HBsAg clearance.

### A clinical trial-enabled study of HBsAg clearance in adults with chronic hepatitis B

To examine mechanisms of HBsAg clearance in CHB patients and make comparisons with the intrahepatic immunity identified in the mouse model, we analyzed longitudinal PBMCs from 31 patients with binary HBsAg outcomes: 30 cases in the BeNEG-DO trial, and an additional patient from a precursor pilot study (see Study Design). Clinical and research blood samples were drawn at baseline (BL, pre-intervention) and at frequent intervals after stopping NAs to monitor safety, HBsAg levels, and to capture clinically relevant events. These timepoints captured initial virologic relapse (iVR), when viremia first became quantifiable after unleashed HBV replication (range 1-5 months), and “peak ALT”, when, within the first 12 months post-NA withdrawal, the anti-HBV cellular immune response resulted in maximal hepatocyte injury assessed by release of alanine aminotransferase (ALT) into blood (Fig. 5A). These serial PBMCs were used to examine dynamic immune composition as well as functional and transcriptional cell states in 15 patients who achieved HBsAg clearance and 16 patients who retained stable HBsAg levels when cytometry by time of flight (CyTOF) and sequencing studies were performed (Fig. 5B-C, table S1). Although average baseline HBsAg levels were lower in the HBsAg clearance group as expected (*34*), patients with similar baseline levels were found in both outcome groups, indicating that HBsAg level may influence but did not solely determine the immune response (Fig. 5B and C, fig. S7A, table S1). Similarly, elevated serum ALT levels, clinically suspected to reflect the strength, as well as the presence, of the HBV-specific immune response, did not associate with outcome (fig. S7B-D), suggesting qualitative response differences. A subset of 12 HBsAg clearance and 12 HBsAg persistence patients were used for CyTOF analysis (Fig. 5D-G) and an overlapping subset of 13 clearance and 11 persistence patients was used for cellular indexing of transcriptomes and epitopes sequencing (CITE-seq) to explore immune composition and dynamics at the single-cell level (Fig. 6).

**Fig. 5:**
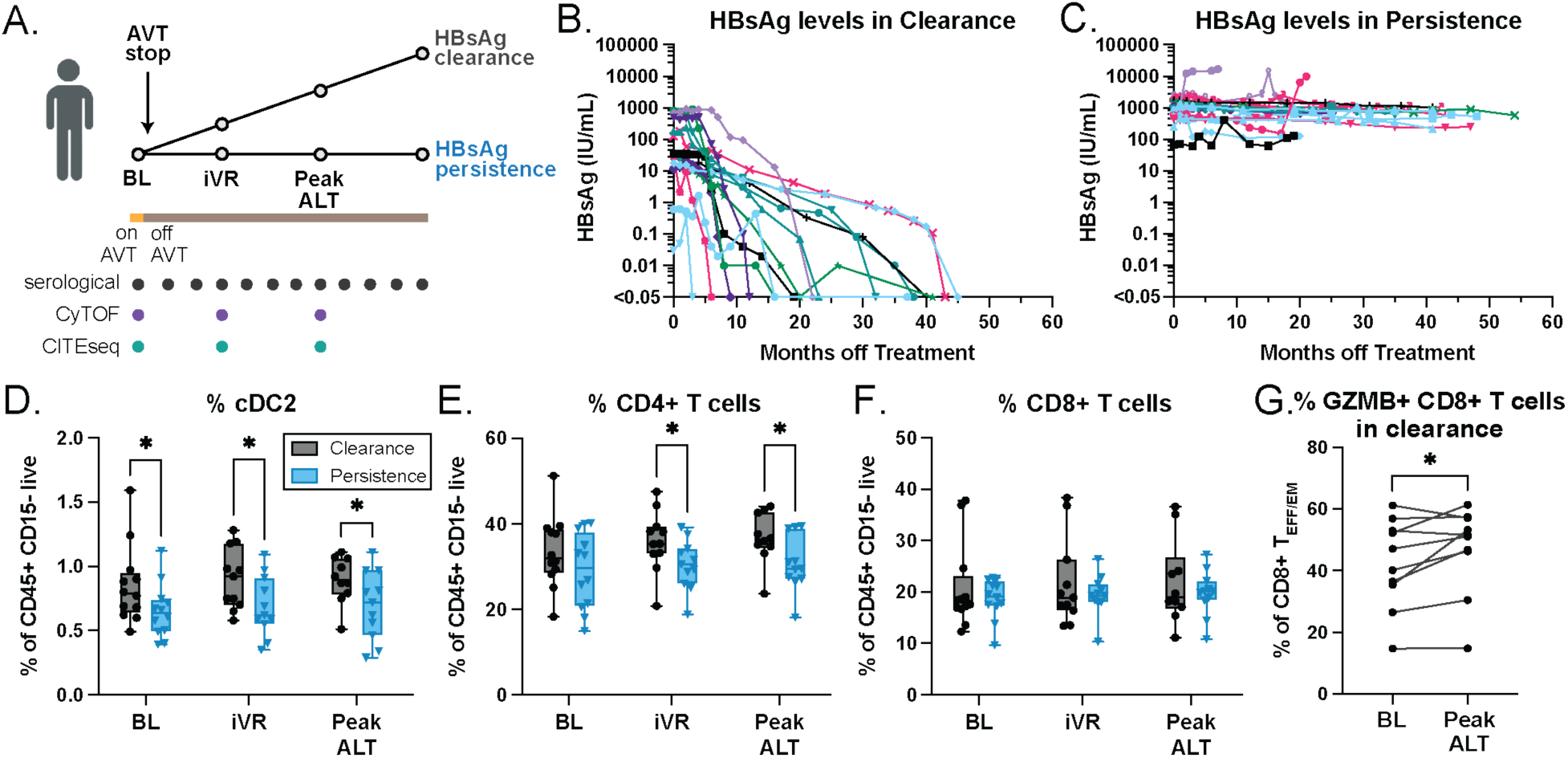
Longitudinal analysis of BeNEG-DO patients with distinct clinical outcomes. (**A**) Schematic depicting longitudinal blood sampling around clinically relevant events from CHB patients after stopping NA. (**B-C**) Quantitative HBsAg kinetics in individually color-coded patients that (**B**) cleared HBsAg (n=15) or (**C**) retained stable levels of HBsAg over time (n=16). (**D**) Percentage of peripheral cDC2s (CD11c+ HLA-DR+ CD16-CD1c+ CD14+/-), (**E**) CD4+ T cells, and (**F**) CD8+ T cells in patients who cleared HBsAg (black) or remained HBsAg+ (blue) as measured by CyTOF. (**G**) Percentage of Granzyme B+ CD8+ T_EFF/EM_ cells (CCR7^-^ CD45RA^-^) at baseline (BL) and Peak ALT in patients who cleared HBsAg. N≥10 per timepoint; statistics performed using (D-F) multiple unpaired t-tests with Benjamini Hochberg FDR correction (*q < 0.1) and (G) paired t-test; * p<0.05.

**Fig. 6:**
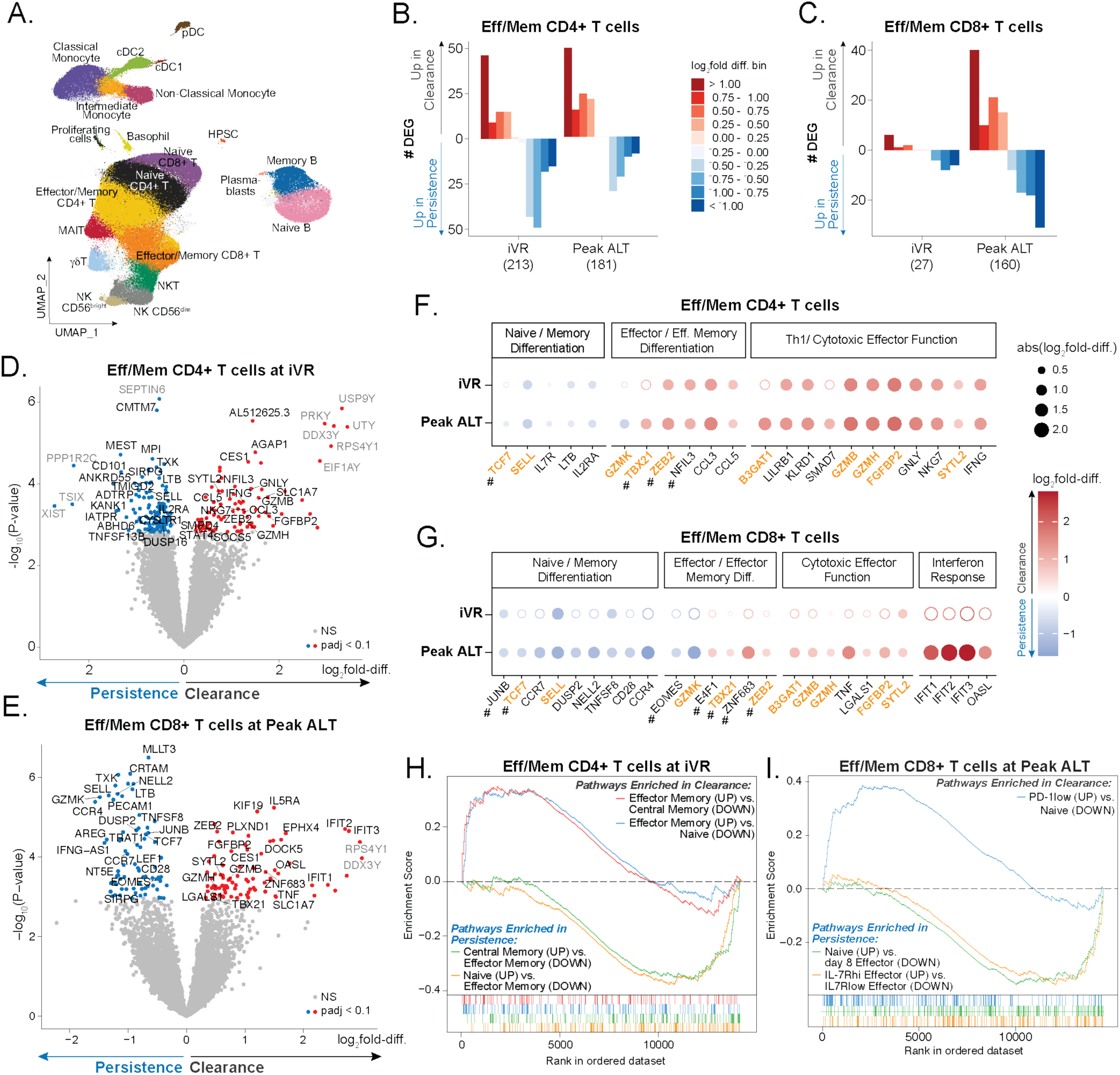
CHB patients who clear HBsAg demonstrate increased transcriptional activity of cytotoxic T cell programs in peripheral blood. Single-cell CITE-seq was performed on PBMC from 24 patients (n=13 HBsAg clearance, n=11 HBsAg persistence) to detect mRNA transcripts and selected proteins. (**A**) Uniform manifold approximation and projection (UMAP) visualization of 519,288 cells clustered by a weighted nearest neighbor (WNN) approach using RNA and protein reads. Number of significant DEG (adj. p<0.1) upregulated in clearance (red) or persistence (blue) patients within (**B**) CD4+ and (**C**) CD8+ effector/memory T cells (all CD4+ and CD8+ T cell clusters excluding naïve clusters) binned by log_2_fold difference; the total number of DEGs is noted in parentheses for each timepoint. Volcano plot depicting DEG in effector/memory (**D**) CD4+ T cells at iVR and (**E**) CD8+ T cells at peak ALT; blue and red dots mark significant DEG (adj. p<0.1) upregulated in persistence and clearance, respectively, with sex-linked genes annotated in gray text. Dot plots depicting DEG (adj. p<0.1, closed circle; NS, open circle) enriched in clearance (red) versus persistence (blue) effector/memory (**F**) CD4+ and (**G**) CD8+ T cells at iVR, and peak ALT, highlighting overlapping transcripts (orange/bold) shared across CD4+ and CD8+ T cell populations and genes that are transcription factors (#). GSEA highlighting significantly enriched pathways from C7 ImmuneSigDB associated with effector/effector memory phenotypes in clearance versus memory/naïve phenotypes in persistence in effector/memory (**H**) CD4+ T cells at iVR and (**I**) CD8+ T cells at peak ALT: GSE3982_EFF_MEMORY_VS_CENT_MEMORY_CD4_TCELL_UP (red, NES = 1.50, adj. p=0.063), GSE26928_NAIVE_VS_EFF_MEMORY_CD4_TCELL_DN (blue, NES=1.49, adj. p=0.049), GSE3982_EFF_MEMORY_VS_CENT_MEMORY_CD4_TCELL_DN (green, NES=-1.59, adj. p=0.022), GSE26928_NAIVE_VS_EFF_MEMORY_CD4_TCELL_UP (orange, NES=-1.69, adj. p=0.004), GSE26495_NAIVE_VS_PD1LOW_CD8_TCELL_DN (blue, NES=1.83, adj. p=0.0007), KAECH_NAIVE_VS_DAY8_EFF_CD8_TCELL_UP (green, NES=-1.50, adj. p=0.047), GSE8678_IL7R_LOW_VS_HIGH_EFF_CD8_TCELL_DN (orange, NES=-1.51, adj. p=0.063).

### Patients who clear HBsAg have more circulating cDC2s and CD4^+^ T cells

Consistent with findings in the mouse model, CyTOF analysis showed that patients who achieved HBsAg clearance exhibited higher abundance of cDC2s at baseline, iVR, and peak ALT, and of CD4^+^ T cells at iVR and peak ALT compared to patients who did not clear HBsAg (Fig. 5D and E). Notably, cDC2s and CD4^+^ T cells were the only cell populations whose frequency differed between HBsAg clearance and persistence groups; all other populations, including other T cell subsets, B cells, DCs, monocytes, and granulocytes were equivalent (fig. S8). Although we did not observe a significant difference in the abundance of CD8^+^ T cells at any timepoint (Fig. 5F), Granzyme B expression in CD8^+^ T cells increased at peak ALT compared to baseline in clearance patients, supporting that activation of a CD8^+^ T cell cytotoxicity program occurs in those who clear HBsAg (Fig. 5G). On the other hand, there was no difference in CD8^+^ T cell expression of markers of exhaustion and memory between clearance and persistence patients (fig. S9), which again parallels our findings in the memory phase of the mouse model (fig. S5D, F). These findings, including the temporal sequence, suggest that CD4^+^ T cell activation and differentiation could play a role in HBsAg clearance in CHB patients, including functional augmentation of the CD8^+^ T cell response shown in the mouse model.

### Single-cell transcriptomic analysis of immune cells in patients with divergent HBsAg outcomes

To profile immune cell subsets and their functional states across timepoints, we performed CITE-seq analysis of PBMCs from 13 patients who achieved HBsAg clearance and 11 with stable HBsAg persistence (fig. S10A). Following the multi-omic integration of high-quality RNA and protein expression and cell filtering that resulted in 519,288 cells, we performed unsupervised clustering and identified 21 main myeloid and lymphoid cell types, including naïve and effector/memory T cells, and defined 34 finer populations representing different cell subsets based on the expression of canonical markers (Fig. 6A, fig. S10B and C, supplemental materials and methods). While most immune cell subsets were equivalently abundant in clearance and persistence groups across all timepoints, patients with HBsAg persistence had higher baseline frequency of naïve CD8^+^ T cells whereas clearance patients demonstrated higher frequency of a CD8^+^ T_EFF_ cell cluster that had high ribosomal gene expression (fig. S10D-F). Notably, a cluster of CD4^+^ CTLs was enriched in HBsAg clearance patients at peak ALT during a strongly active phase of the adaptive response (fig. S10D and G), a cell subset associated with HBsAg clearance in the mouse model (Fig. 1L and M, fig. S1I-L).

To investigate gene expression differences associated with HBsAg clearance and persistence we built a series of linear mixed-effects models (limma (*35*), Methods). This approach allowed us to leverage our longitudinal samples to adjust for patient-specific effects. Balancing this adjustment with interpretability of the longitudinal models, we chose to create models of timepoint pairs: baseline and iVR (“iVR”), and baseline and peak ALT (“peak ALT”). These timepoint pair models were constructed to compare differential gene expression in HBsAg clearance versus persistence groups within each cell population. This analysis revealed the strongest signal was primarily observed when modeling the main immune cell types, with very few significant differences observed for the finer immune cell subsets, likely due to those populations having fewer numbers of cells.

Since the mouse model strongly implicated effector T cell subsets in driving HBsAg clearance, we focused our analysis on the transcriptional differences between clearance and persistence in non-naïve, effector/memory CD4^+^ and CD8^+^ T cells (i.e., all T cell clusters excluding naïve clusters). Comparative analysis of clearance versus persistence groups revealed numerous differentially expressed genes (DEGs) in effector/memory T cells (Fig. 6B and C, data file S1). Effector/memory CD4^+^ T cells exhibited differential expression of 213 genes at iVR and 181 genes at peak ALT, suggesting substantial outcome-associated transcriptomic changes during the early post-treatment phase. (Fig. 6B). Conversely, in effector/memory CD8^+^ T cells, there were only 27 DEGs at iVR compared to a more robust signal of 160 DEGs at peak ALT (86 DEGs associated with clearance; 74 with persistence), potentially indicating a later peak response to treatment withdrawal (Fig. 6C).

### Patients who clear HBsAg engage overlapping transcriptional programs in effector CD4^+^ and CD8^+^ T cells

Analysis of the DEGs at iVR and peak ALT highlighted that HBsAg outcomes were associated with divergent T cell states throughout the post-treatment response. Notably, the topmost DEGs in both CD4^+^ and CD8^+^ Effector/Memory T cells included transcripts associated with T cell differentiation and effector responses (Fig. 6D-G), mirroring programs that associated with HBsAg outcome in the mouse model. Specifically, CD4^+^ T cells in clearance patients upregulated transcripts associated with T_H_1 differentiation including transcription factors (*TBX21*, *STAT4*, *ZEB2*, *NFIL3, SOCS5*), cytokines (*IFNG*), and chemokines (*CCL3*, *CCL5*). Furthermore, CD4^+^ T cells associated with HBsAg clearance also demonstrated upregulation of markers associated with effector differentiation and cytotoxic function (*GZMB*, *GZMH*, *B3GAT1*, *FGFBP2*, *GNLY*, *NKG7*, and *SYTL2*). Consistent with the effector CD4^+^ T cell transcriptional programs and the CyTOF data (Fig. 5G), CD8^+^ T cells from HBsAg clearance patients were enriched in many of the same transcripts associated with T_H_1 and effector CTL responses, in addition to enrichment of transcripts involved in the interferon response (*IFIT1*, *IFIT2*, *IFIT3*, *OASL*).

In contrast to the strong effector T cell signature observed in HBsAg clearance, CD4^+^ and CD8^+^ T cells from patients with HBsAg persistence demonstrated upregulation of genes associated with a naïve, quiescent, or less activated and differentiated state. Specifically, CD4^+^ T cells from persistence patients had higher levels of multiple transcription factors and markers associated with central memory and stemness (*TCF7*, *SELL*, *IL7R*). In addition to memory-associated transcripts, some in common with the CD4^+^ T cells (Fig. 6G), CD8^+^ T cells in persistence patients upregulated additional markers of central memory (*CCR7, CD28, TNFSF8, LEF1*), and notably showed enriched expression of *EOMES* and *GZMK*, consistent with a profile of dysfunctional CD8^+^ T cells associated with pathologic inflammation (*36*). This HBsAg persistence transcriptomic profile contrasted with the *GZMB-* and *GZMH-*enriched CD8^+^ T cells observed in clearance, markers typically associated with effective cellular immunity. CD4^+^ T cells in patients with HBsAg persistence were also enriched for transcripts associated with Th2 polarity (*CYSLTR1*, *DUSP16*) and activation-limiting potential (*CD101*, *SIRPG*, *TMIGD2*). Notably, the CD8^+^ T cells in persistence were similarly enriched for transcripts offering activation limiting potential (*PECAM1, CMC1*) as well as an antisense RNA (*IFNG-AS1*) associated with IFN-γ suppression in memory T cells (*37*).

Gene set enrichment analysis (GSEA) (*38, 39*) on the differential gene expression results corroborated our observation of a strong dichotomy between a distinctly activated effector state in patients achieving HBsAg clearance versus a less differentiated “naïve” state in patients with HBsAg persistence (data file S2). Specifically, pathways associated with a more stem-like or central memory phenotype were enriched within CD4^+^ T cells from patients with HBsAg persistence while an effector-memory pathway was enriched in CD4^+^ T cells from clearance patients (Fig. 6H) (*40*). Similarly, naïve and memory pathways were enriched in CD8^+^ T cells from HBsAg persistence patients, in comparison to the effector pathways enriched in patients achieving HBsAg clearance (Fig. 6I). In aggregate, this transcriptomic analysis therefore identified distinct CD4^+^ and CD8^+^ T cell differentiation programs associated with HBsAg clearance that closely align with the programs identified in the mouse model.

## DISCUSSION

Our findings in the mouse model and in patients with CHB identify parallel effector immune mechanisms that lead to HBsAg clearance. Upstream of the T cell responses studied in both settings, the mouse model allowed us to demonstrate a pivotal role for hepatic dendritic cells, particularly cDC2s, in directing effective HBV-specific CD4^+^ T cell activation and differentiation, which we and others have shown to be crucial for HBsAg clearance. In young mice, we identify a major defect in the ability of resident hepatic DCs to activate CD4^+^ T cells, and using MHCII tetramers, we show that this defect results in a short-lived, ‘failure-to-thrive’ phenotype of HBsAg-specific CD4^+^ T cells. In addition, we demonstrate that hepatic lymphoid organization and effective HBsAg-specific CD8^+^ T cell function are critically dependent on CD4^+^ T cell help. Notably, hepatic immune composition and phenotyping revealed that cDC2s, and to a lesser extent cDC1s, are less abundant and show continued signs of functional impairment as the young HBsAg^+^ animals became adult “chronic” mice, and also that HBV-specific CD4^+^ T cells were not detectable in their livers. In contrast, HBV-specific CD8^+^ T cell abundance was equivalent and robust in these chronic mice, without clear evidence of differential exhaustion. Together, these data provided a mechanistic window into the ineffective immunity that develops in the young liver leading to HBV persistence, and a prediction that the hepatic immune response in CHB, which is largely a consequence of infection at a young age, includes less effective DCs and impaired CD4^+^ T cell activation and memory.

Longitudinal samples from the BeNEG-DO clinical trial offered rare opportunity to analyze the developing immune response that leads to HBsAg clearance or continued persistence in patients with CHB after stopping NA therapy, and to probe for evidence of shared mechanisms identified in the mouse model. While we could not directly study longitudinal hepatic immune composition and phenotypes in our patients, we were able to study immune composition in longitudinal PBMCs, sampling key transitions and outcomes. These studies revealed striking mechanistic congruence. Our CyTOF analysis revealed that HBsAg clearance was distinguished by higher abundance of cDC2s at baseline and all subsequent timepoints, and increased CD4^+^ T cell abundance at iVR and peak ALT. Although CD8^+^ T cell abundance was similar at each timepoint in each HBsAg outcome group, as in the mouse model, CD8^+^ T cells showed evidence of increased effector function over time, which we consider a likely consequence of CD4^+^ T cell help.

Single-cell sequencing analysis of patient samples revealed outcome-associated transcriptional activity among effector/memory CD4^+^ T cells at iVR and peak ALT, identifying transcripts associated with distinct cell subsets and states in each HBsAg outcome. At iVR and continuing at peak ALT, HBsAg clearance patients exhibited transcriptional programs canonically associated with T_H_1 differentiation, an effector memory state, and signature transcripts expressed by cytotoxic cells. This finding of cytotoxic CD4^+^ T cell activation in a population of cells obtained *ex vivo*, as in the mice, and its association with HBsAg clearance is further support for an active effector role for these cells in viral control, as proposed 33 years ago (*41*) and more recently by others (*42*). In patients with HBsAg persistence, the DEGs identified in CD4^+^ T cells at iVR and peak ALT included transcripts associated with central memory and a less differentiated state. These findings implicated strong effector CD4^+^ T cell differentiation and activation as a distinguishing feature of HBsAg clearance and are consistent with our data in the mouse model, and other studies (*15, 16, 43, 44*).

Contrasting with the CD4^+^ T cells, single-cell analysis of effector/memory CD8^+^ T cells found relatively few DEGs at iVR, with robust outcome-associated differences apparent only when considering peak ALT. In HBsAg clearance, the CD8^+^ T cells upregulated signature transcripts associated with effector function, including cytotoxicity (consistent with CyTOF data) and a type I interferon response, whereas in HBsAg persistence, the CD8^+^ T cells upregulated transcripts characteristic of a central memory, naïve-like state and an inflammatory, non-cytolytic phenotype. Importantly, when taken together with T cell dynamics observed during HBsAg clearance in mice, the high numbers of differentially expressed genes in CD8^+^ T cells observed at peak ALT but not at iVR – when CD4^+^ T cells had many DEGs – suggests that the CD4^+^ T cell response may be preceding and augmenting the CD8^+^ T cell response.

After CHB patients unleash HBV replication by stopping NAs after long-term suppression, the immune mechanisms of HBsAg clearance invoked by our data, supported by the mouse model, are reminiscent of those in acute HBV infection. In contrast to the dysfunctional, inflammatory T cell response in CHB that originally necessitates NA therapy, patients achieving HBsAg clearance after stopping NA therapy appear to engage an immune response akin to an infection-terminating response in acutely infected adults and in adult mice in our model. On the other hand, the T cell response in our patients with HBsAg persistence likely recapitulated the original program that led to CHB, a program we now suspect was initiated with minimal assistance from HBV-specific CD4^+^ T cells. While studies of the hepatic mechanisms responsible for the differential T cell responses were not tractable in our patients, the mouse model did provide an experimental gateway to this avenue of investigation, which holds therapeutic potential.

Our data shifts the immunological paradigms of HBsAg clearance and persistence to provide a focus on dendritic cells and their ability to appropriately sense HBV, and prime and perpetuate an HBV-specific CD4^+^ T cell response. We propose a model in which parallel immune mechanisms can induce HBsAg clearance in both acute and chronic HBV infection. In this model, an effective CD4^+^ T cell response governs further DC licensing, hepatic lymphoid organization, as well as augmenting HBV-specific CD8^+^ T cell and B cell effector function, leading to the orchestrated immune response that ultimately controls the infection. Our data also suggest a role for cytotoxic CD4^+^ T cells in this process. The CHB patients who went on to clear HBsAg had signs at baseline that this effective immunological program had already been partially engaged, opening the possibility that predictive proteomic/genomic algorithms could be developed to select patients who would benefit from treatment withdrawal. This new data also identifies rational therapeutic avenues to terminate CHB including targeting CD4^+^ T cell co-stimulatory pathways and DC activation to increase HBV-specific CD4^+^ T cell abundance, effector function, and hepatic positioning. Such therapeutics could be used in conjunction with timed treatment withdrawal or as stand-alone approaches.

## MATERIALS AND METHODS

### Study design

This human and animal model study was designed to identify immune mechanism(s) of HBsAg clearance and persistence. For the model, all mice were bred and housed in specific pathogen-free housing under an IACUC approved protocol (AN170936) and in accordance with the guidelines of the Laboratory Animal Resource Center of the University of California, San Francisco. We used male and female mice in all experiments. WT C57BL/6 mice were purchased from Jackson Laboratory then bred at UCSF. For the mouse experiments, young (3-3.5 weeks old, pre-weaning) or adult (>8 weeks old) HBVtg*Rag1*^-/-^ mice (*14*) were given 0.8-1.0×10^8^ syngeneic splenocytes pooled from adult (>8 weeks old) WT mice via tail vein injection. Mice were bled at regular intervals on days 5, 7, 14, 21, 28, 56, and 84, unless otherwise specified, and followed for plasma ALT, HBsAg and anti-HBs. Hepatic leukocytes and/or whole liver tissue pieces were collected at pre-defined endpoints on days 3, 5, 8, 15, 30, and 60-120 post-adoptive transfer for further characterization by spectral flow cytometry, immunohistochemistry, and *in vitro*functional assays.

Our mouse studies featured several comparisons of immunologic parameters between adult and young and/or in CD4^+^ T cell-depleted mice. A minimum of 5-8 mice per group were used for these comparisons, which provided adequate power to identify differences between comparators in our published work using similar numbers of animals (*15–17*). The number of animals used, replicate experiments, and specific statistical tests applied are detailed in the figure legends.

Of the 31 patients studied here, 30 were enrolled as cases in the two-center BeNEG-DO clinical trial (https://clinicaltrials.gov/study/NCT02845401) and 1 case in our precursor “BeNEG” pilot study. BeNEG-DO is a prospective case-control study of safety and clinical outcomes in adult human subjects with e antigen negative CHB who either stop or continue NA antiviral therapy. For this immunological study, CyTOF and sequencing studies were performed using PBMCs from cases with the stated binary HBsAg outcomes who had provided adequate serial timepoint samples for this research. Before stopping NA, all cases: (i) were serologically HBsAg positive, anti-HBs negative, e-antigen negative, and anti-HBe positive; (ii) had immune-active e antigen negative CHB when NAs were started; (iii) had completed ≥192 weeks of NA therapy and maintained complete virologic responses (serum HBV DNA <20 IU/ml); (iv) at end of treatment, none had advanced hepatic fibrosis as determined by biopsy (≤F2) and non-invasive tests, and none had significant additional liver disease, co-infection, or co-morbidities. The two outcome groups were not distinguished demographically or by HBV genotype (Table S1). Informed written consent was obtained from all study subjects. Longitudinal blood samples were obtained under IRB protocol #11-07994 at UCSF and Sutter Health (BeNEG-DO) and protocol 2012.059-2 (SCoo) at Sutter Health (BeNEG). Synchronous clinical and research blood samples were collected monthly for the first 6 months after NA withdrawal, every 2 months during months 7-12, and every 3 months beyond month 12. Lab monitoring frequency exceeded protocol during ALT flares (ALT≥200 U/L) and/or at the discretion of the managing hepatologist. Re-treatment criteria, including for cases with immune active disease without virologic control, can be found in the clinical trial description, NCT02845401. A healthy donor sample purchased from Vitalant Research Institute (San Francisco, CA) was used as a batch control for CyTOF experiments. All clinical tests were performed at local CLIA-approved laboratories unless otherwise indicated. All research samples were centrally processed and stored in the Ibrahim El-Hefni Liver Biorepository. Quantitative HBsAg (qHBsAg) was measured on an Abbott ARCHITECT instrument per the manufacturer’s instructions.

### Statistical analysis

Mouse model: Unless otherwise specified, statistics were performed using Prism (Graph Pad Software). Statistical significance was determined by two-tailed unpaired Student’s *t* test, two-tailed paired Student’s *t* test (for paired data on the same mice), one-way ANOVA with Dunnett’s correction for multiple comparisons to a control group, or one-way ANOVA with Tukey’s multiple-comparison test (when multiple groups were all compared to each other), or two-way ANOVA with Dunnett’s correction for multiple comparisons to a control group (when multiple parameters were compared between more than two groups and a defined control group), or two-way ANOVA with Tukey’s multiple-comparison test (when one parameter was compared for more than two biological groups, and all groups were compared to each other). For significance of HBsAg clearance, in which one of two possible outcomes was compared (clearance=1 versus no clearance=0), log-rank (Mantel-Cox) Chi-square test was used. For ordinal or ranked data (histology scores), the Mann-Whitney rank sum test was used. In all figures with multiple *n*, data are presented as means ± SEM; p < 0.05 was considered significant.

Human studies: Statistical analysis of CyTOF was performed using GraphPad Prism (v10.4.1) and CITE-seq using a suite of R packages (v4.1.1) as described in Supplementary Materials and Methods. The p-values were adjusted for multiple tests using Benjamini-Hochberg procedure (BH), and the adjusted p-values (adj. p) were used to determine statistical significance. CyTOF frequencies were compared between groups using unpaired t-tests with adj. p<0.05. Normalized cell frequencies from CITE-seq data were compared between groups of interest using Wilcoxon Rank-Sum test with adj. p<0.1. We used the voom function of the limma package (v3.48.3) to assess differential expression, computing the log-odds of differential expression with attendant empirical Bayes moderation of the standard errors and considered genes with adj. p<0.1 as significant. Pathway enrichment analysis was performed using gene set enrichment analysis (GSEA) with adj. p<0.1.

## Supporting information

DEG in CD4+ effector_memory T cells at iVR

DEG in CD4+ effector_memory T cells at peak ALT

DEG in CD8+ effector_memory T cells at iVR

DEG in CD8+ effector_memory T cells at peak ALT

GSEA in CD4+ effector_memory T cells at iVR

GSEA in CD8+ effector_memory T cells at peak ALT

TotalSeq-A antibody panel

## List of Supplementary Materials

Fig. S1: Young mice fail to clear HBsAg effectively and exhibit defective CD4+ T cell responses.

Fig. S2: Representative gating demonstrating adult CD4+ T cells express higher levels of IFNγ, CXCR6, Tbet, CX_3_CR1, Granzyme B, and Perforin compared to young CD4+ T cells.

Fig. S3: Age-dependent differences in cDC populations are also apparent in wild-type mice and are derived from recipient origin.

Fig. S4: Splenic APCs from young mice are not impaired in their ability to activated naïve CD4+ T cells

Fig. S5: Young mice and CD4-depleted mice exhibit impaired CD8+ T cell function accompanied by a decrease in cDC activity.

Fig. S6: Representative gating demonstrating that CD8+ T cells in adult mice express higher levels of IFNγ, Granzyme B, and Perforin compared to CD8+ T cells in young mice.

Fig. S7: Baseline HBsAg levels and dynamics of ALT elevation in study patients.

Fig. S8: Frequencies of immune cell subsets that are similar in HBsAg clearance and persistence patients.

Fig. S9: Peripheral CD8+ T cells from CHB patients that clear or retain circulating HBsAg do not differ in their expression of canonical “exhaustion/dysfunction” markers.

Fig. S10: CITE-seq sample collection, single cell population identity, and population frequency.

Table S1: Characteristics of CHB patients who stopped NA therapy.

## Supplementary Materials and Methods

Data file S1: DEG in CD4+ effector_memory T cells at iVR.

Data file S2: DEG in CD4+ effector_memory T cells at peak ALT.

Data file S3: DEG in CD8+ effector_memory T cells at iVR.

Data file S4: DEG in CD8+ effector_memory T cells at peak ALT.

Data file S5: GSEA in CD4+ effector_memory T cells at iVR.

Data file S6: GSEA in CD8+ effector_memory T cells at peak ALT.

Data file S7: TotalSeq-A antibody panel.

## Acknowledgments

We are indebted to all of the patients who participated in our studies as well as the contributing clinicians: Erick P. Chan, Eddie Cheung, Edward W. Holt, Mary Hu, Jennifer Lai, Tammy Lee, Jiayi Li, Neil Mehta, Raphael Merriman, Sara Miller, Marion Peters, Jennifer Roost, Brennan Scott, Norah Terrault, Marice Thomas, Francis Yao, Kidist Yimam, Hans Yu. We thank Dan Bunis for data processing, Tiffany Yeh for HBV genotyping, and David Stone, Stephanie Huynh and Diana Suen for biosample processing. We also thank the BeNEG-DO Data Safety Monitoring Board for overseeing safety and data quality. We acknowledge support from the Ibrahim El-Hefni Liver Biorepository, UCSF Liver Center (NIH P30 DK026743), the California Liver Institute, the Norman Raab Foundation, the Technical Training Foundation, the Ing Foundation, and the UCSF Data Science CoLab, Disease to Biology CoLab, and UCSF Parnassus Flow Cytometry CoLab (RRID:SCR_018206) supported in part by Grant NIH P30 DK063720 and by the NIH S10 Instrumentation Grant 1S10OD026940-01. We thank the NIH Tetramer Core Facility (NIH Contract 75N93020D00005 and RRID:SCR_026557) for providing the MHCII and MHCI tetramers loaded with HBsAg_126-138_, HBsAg_321-335_, HBsAg_353-360_, and HBsAg_370-378_ peptides.

## Author contributions

Conceptualization: JLB, SC, GKF, JJ, LA, JP, AEW

Methodology: JMJ, LA, JP, SC, JLB, NDC, AWE, SS, NC, NW, AE, ST, AR, CPL, ST, MS (Stec), MA, GC, JS, JL, MS (Simone), MS (Sarkar)

Investigation: SC, JLB, GKF, JMJ, LA, JP, NDC, RKP, AE, MA, MS, AJC, MRS

Visualization: RKP, AWE, LA, JMJ, JP, NDC

Funding acquisition: JLB, SC, AEW

Project administration: JLB, SC, GKF

Supervision: JLB, SC, GKF, AJC, MRS

Writing – original draft: JLB, JMJ, SC

Writing – review & editing: JLB, JMJ, SC, GKF, NDC, LA, RKP

## Data and materials availability

All data and animal models are available upon request. CITE-seq data of the CHB trial patients is available at GEO: GSE291286.

