## Supplemental Figures and Methods for "Clinical cure of chronic hepatitis B is dependent on activation and perpetuation of robust CD4^+^ T cell responses"

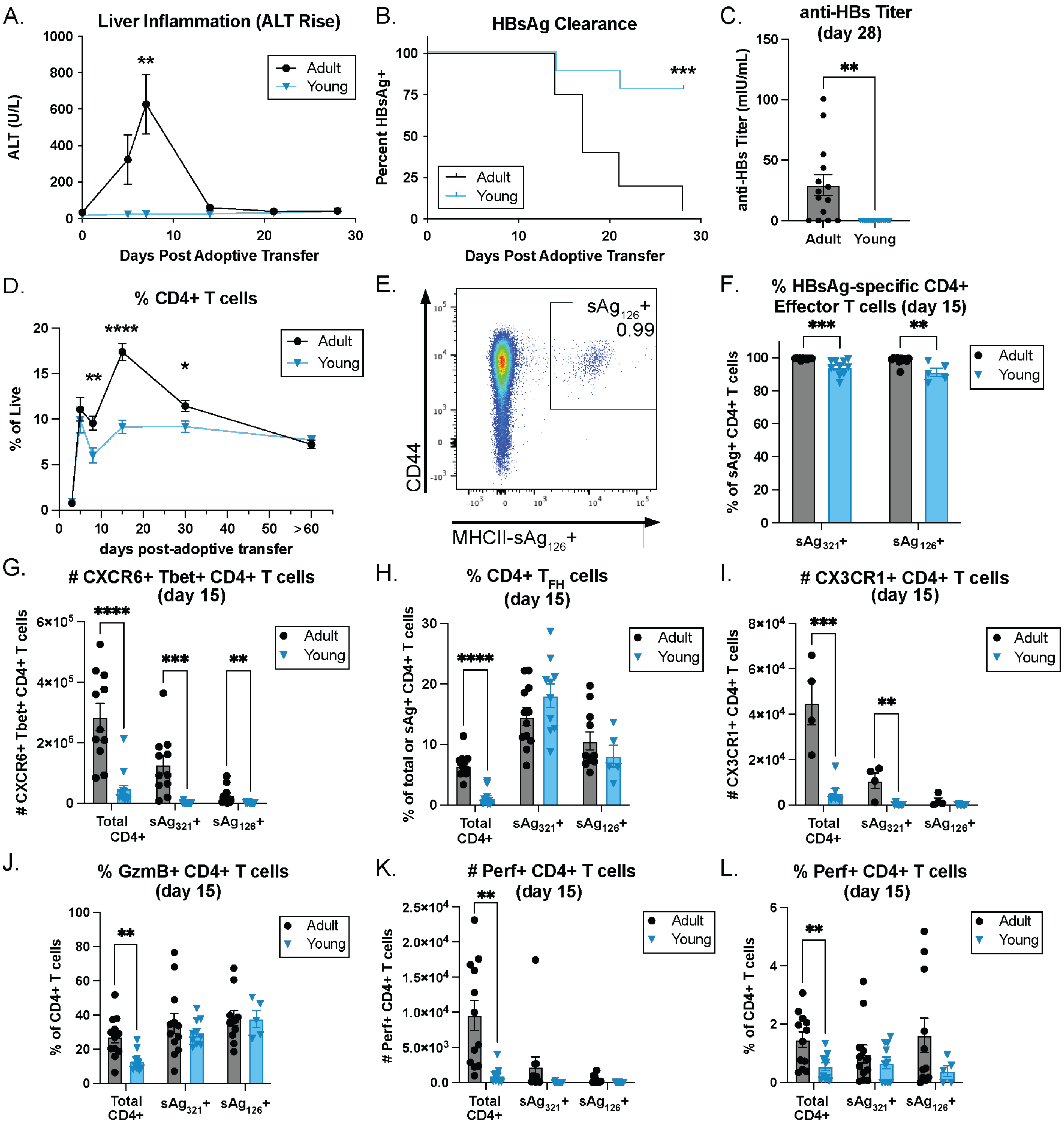


**Fig. S1: Young mice fail to clear HBsAg effectively and exhibit defective CD4+ T cell responses.** Adult (black circles) and young (blue triangles) HBVtgRag$^{-/-}$ mice were bled at the indicated timepoints post-adoptive transfer of WT splenocytes and assayed for (**A**) ALT, (**B**) HBsAg, and (**C**) HBsAb titers. Hepatic leukocytes were isolated from these mice at the indicated time points and assessed by flow cytometry for (**D**) percentage of total CD4+ T cells. (**E**) Representative flow plot demonstrating staining with the sAg$_{126}$ tetramer. (**F**) Percentage of CD44+ CD62L+ effector sAg$_{321}$- and sAg$_{126}$-specific CD4+ T cells. Number of (**G**) CXCR6+ Tbet+, (**I**) CX3CR1+, and (**K**) Perf+ total and HBsAg-specific CD4+ T cells. Percentage of (**H**) CXCR5+ PD-1+ T$_{FH}$, (**J**) GzmB+, and (**L**) Perf+ total and HBsAg-specific CD4+ T cells. N≥11; statistics performed using (A, C-N) unpaired t-test without multiple test correction or (B) log-rank test.

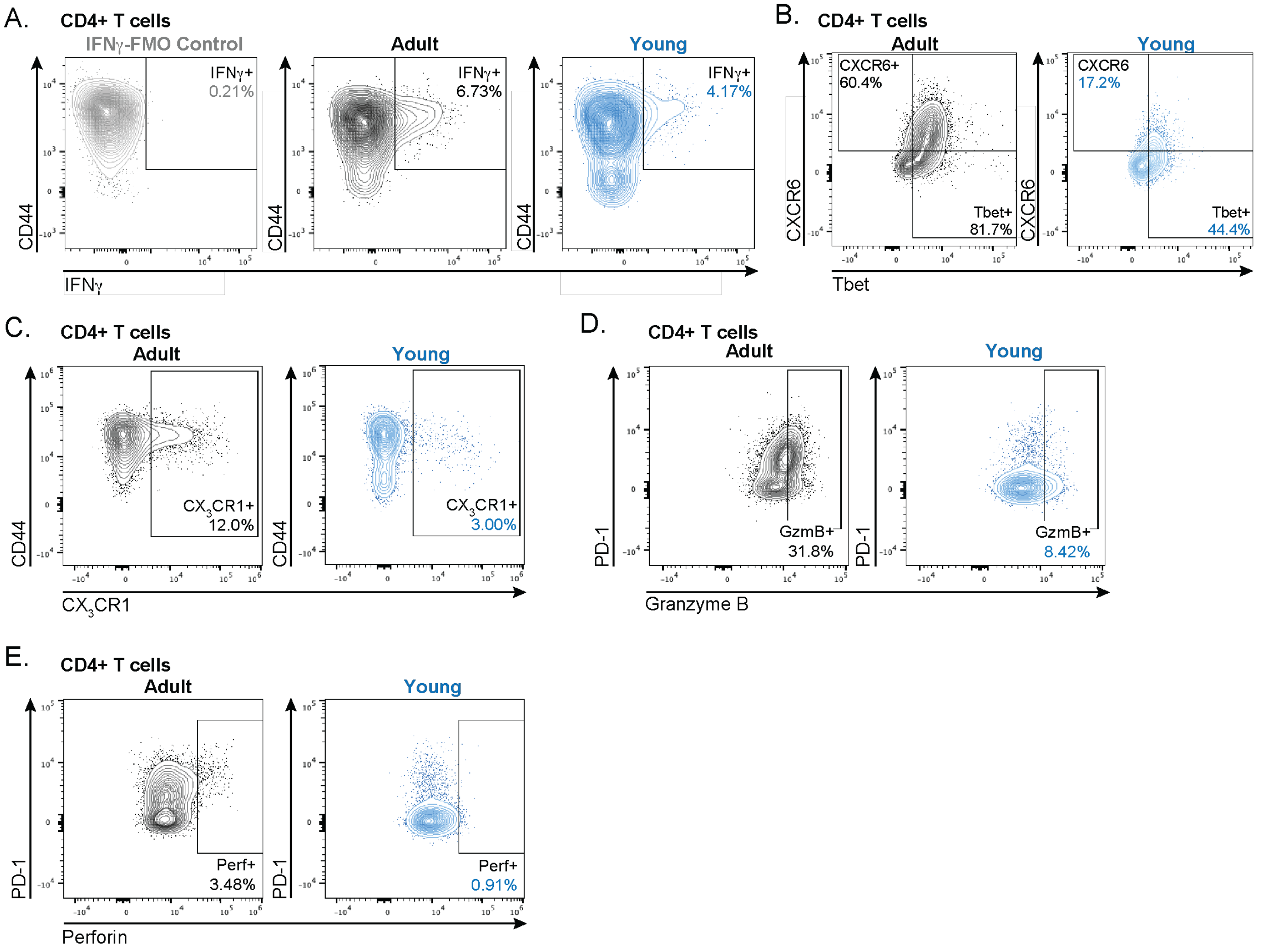


**Fig. S2: Representative gating demonstrating adult CD4+ T cells express higher levels of IFNγ, CXCR6, Tbet, CX3CR1, Granzyme B, and Perforin compared to young CD4+ T cells.** Representative staining of adult and young (**A**) IFNγ+ CD4+ T cells on day 8 and (**B**) CXCR6+ Tbet+, (**C**) $CX_3CR1$+, (**D**) Granzyme B+, and (**E**) Perforin+ CD4+ T cells on day 15 post-adoptive transfer.

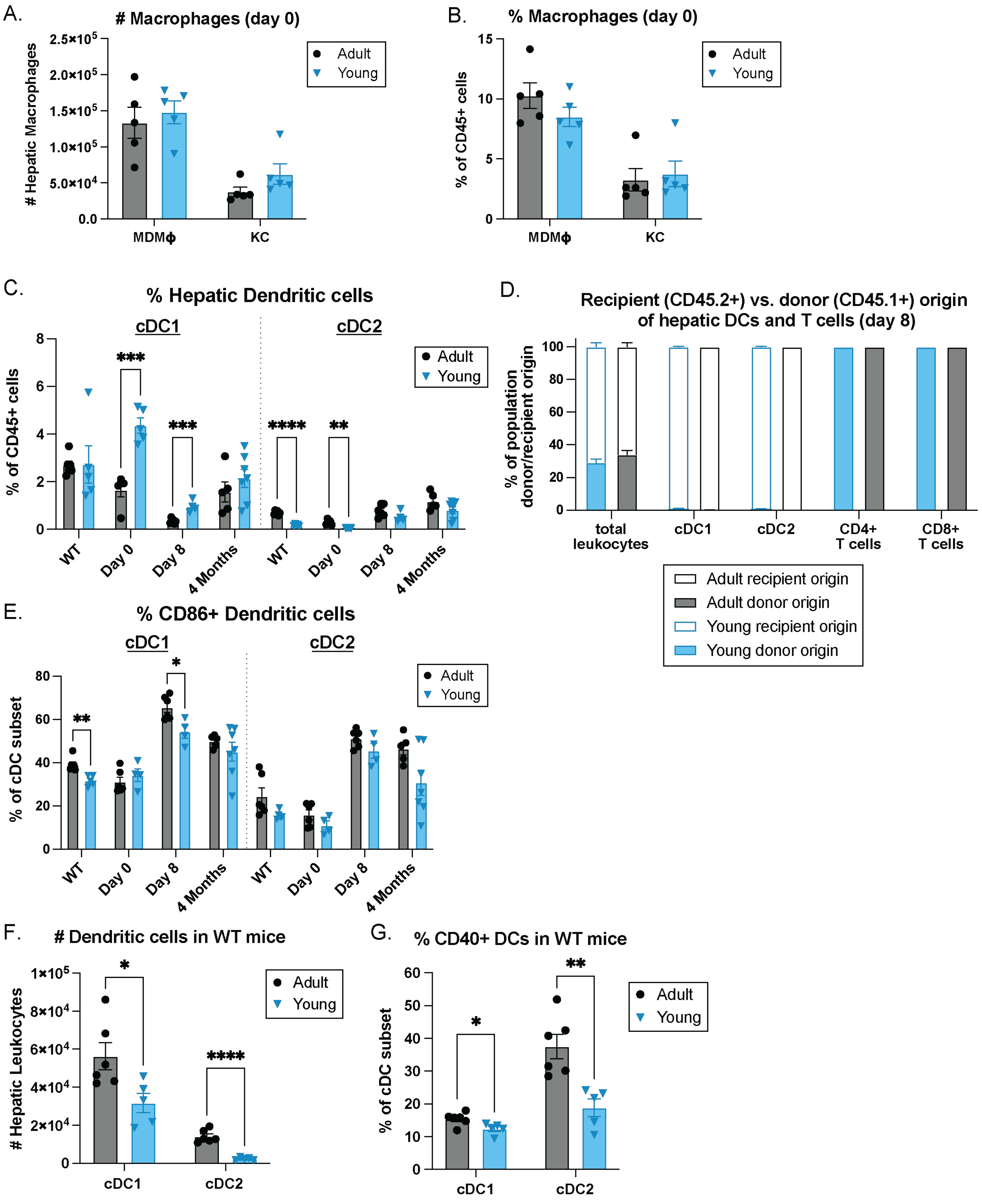


**Fig. S3: Age-dependent differences in cDC populations are also apparent in wild-type mice and are derived from recipient origin.** Hepatic leukocytes were isolated from adult (black circles) and young (blue triangles) HBVtgRag-/- mice prior to adoptive transfer (Day 0) or at the specified timepoints post-transfer, as well as from WT mice. Number (**A**) and percentage (**B**) of MDMϕ (NK1.1- Ly6g- F4/80$^{+/int}$ CD11b$^{hi}$ CD11c$^{–/lo}$) and Kupffer cells (NK1.1- Ly6g- F4/80$^{hi}$ CD11b$^{int}$ CD11c$^{–/lo}$ CD64+). (**C**) Percentage of cDC1 (NK1.1- Ly6G- F4/80- CD11b- MHCII+ CD11c+ XCR1+ CD172a-) and cDC2 (NK1.1- Ly6G- F4/80- MHCII+ CD11c+ XCR1- CD172a+). (**D**) Adult and young HBVtgRag-/- mice (CD45.2+) were adoptively transferred with CD45.1+ splenocytes to differentiate between donor (CD45.1+) and recipient (CD45.2+) origin, and the fraction of leukocytes, cDCs, CD4+, and CD8+ T cells from each origin were determined (N≥4). (**E**) Percentage of CD86+ cDCs. (**F**) Number of Hepatic cDC1 and cDC2 from adult and young WT mice. (**G**) Percentage of CD40+ cDCs in adult and young WT mice. (A-E) N≥4 (F-G) N≥6**;** statistics performed using unpaired t-test without multiple test correction.

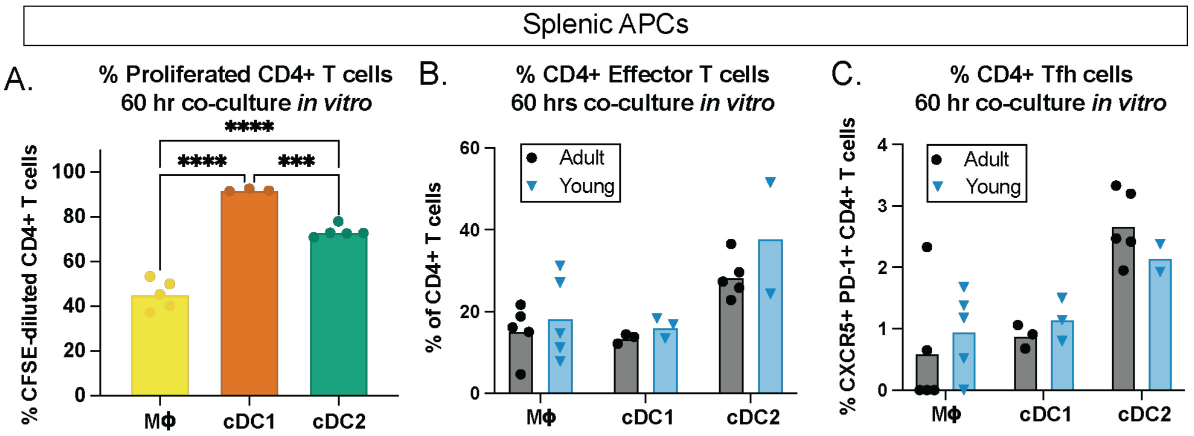


**Fig. S4: Splenic APCs from young mice are not impaired in their ability to activate naïve CD4+ T cells.** Splenocytes were isolated from adult (black circles) and young (blue triangles) HBVtgRag-/- mice and sorted by FACS for macrophages (Mϕ; CD45+ NK1.1- CD90.2- MHCII+ CD11c - F4/80+ CD64+ CD11b+); cDC1 (CD45+ NK1.1- CD90.2- F4/80- CD64- MHCII+ CD11c+ XCR1+ CD172a-); and cDC2; (CD45+ NK1.1- CD90.2- F4/80- CD64- MHCII+ CD11c+ XCR1- CD172a+). Splenic APCs were co-cultured with CD4+ OT-II lymphocytes in the presence $Ova_{323\text{-}339}$ peptide for 60 hours and subsequently assessed by flow cytometry. (**A**) Percentage of CD4+ T cells that underwent proliferation upon stimulation with adult APCs. Percentage of CD4+ T cells with an (**B**) effector phenotype (CD44+ CD62L-) and (**C**) T follicular cell ($T_{FH}$) phenotype (CXCR5+ PD1+). N=pooled from ≥8 mice, dots represent technical replicates; statistics performed using unpaired t-test without multiple test correction.

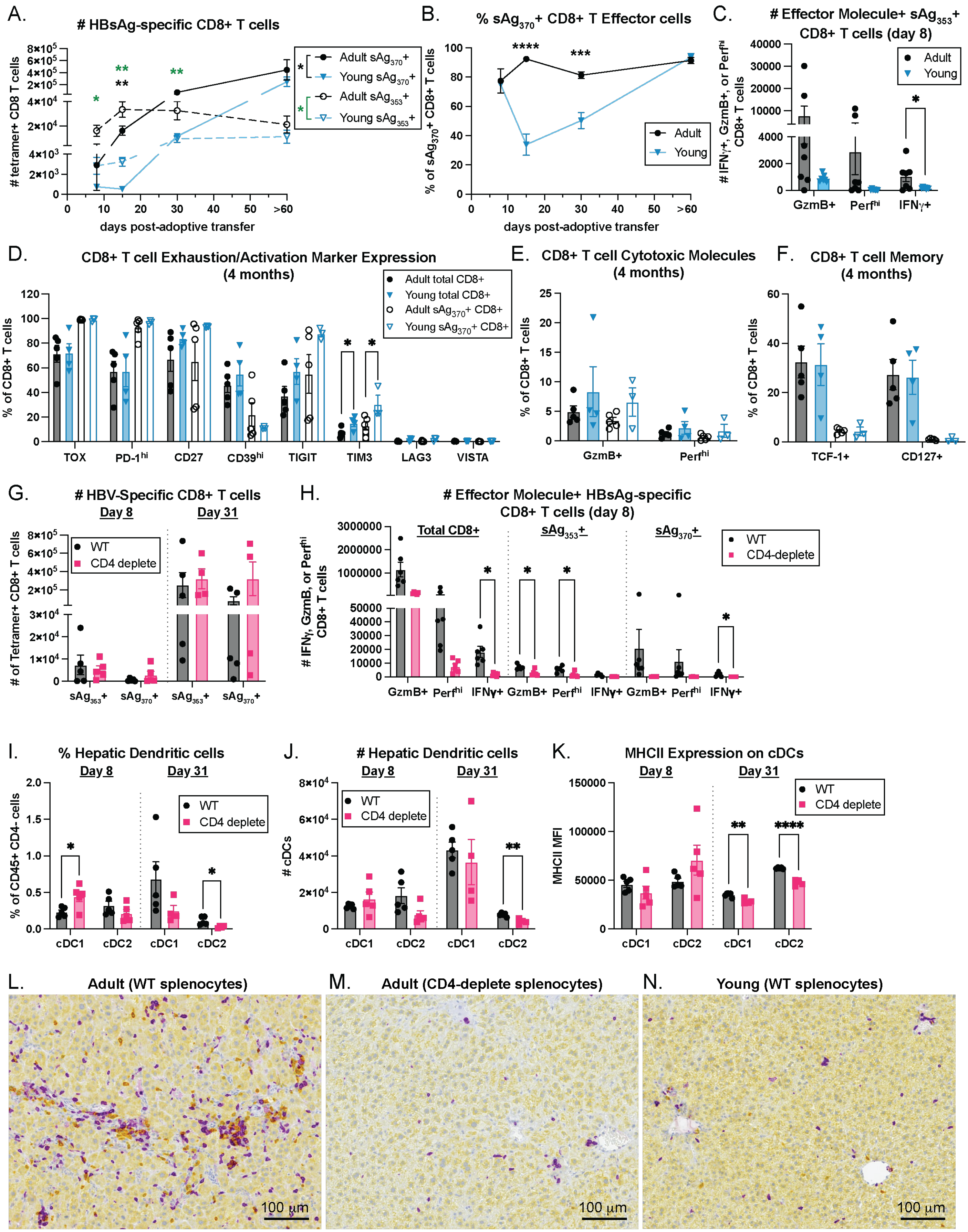

A.
### HBsAg-specific CD8+ T cells
### tetramer+ CD8 T cells
days post-adoptive transfer
Adult sAg370+
Young sAg370+
Adult sAg353+
Young sAg353+
B.
% sAg370+ CD8+ T Effector cells
% of sAg370+ CD8+ T cells
days post-adoptive transfer
Adult
Young
C.
### Effector Molecule+ sAg353+ CD8+ T cells (day 8)
### IFNγ+, GzmB+, or Perfhi CD8+ T cells
Adult
Young
GzmB+
Perfhi
IFNγ+
D.
CD8+ T cell Exhaustion/Activation Marker Expression (4 months)
% of CD8+ T cells
Adult total CD8+
Young total CD8+
Adult sAg370+ CD8+
Young sAg370+ CD8+
TOX
PD-1hi
CD27
CD39hi
TIGIT
TIM3
LAG3
VISTA
E.
CD8+ T cell Cytotoxic Molecules (4 months)
% of CD8+ T cells
GzmB+
Perfhi
F.
CD8+ T cell Memory (4 months)
% of CD8+ T cells
TCF-1+
CD127+
G.
### HBV-Specific CD8+ T cells
Day 8
Day 31
### of Tetramer+ CD8+ T cells
WT
CD4 deplete
sAg353+
sAg370+
H.
### Effector Molecule+ HBsAg-specific CD8+ T cells (day 8)
### IFNγ, GzmB, or Perfhi CD8+ T cells
Total CD8+
sAg353+
sAg370+
WT
CD4-deplete
GzmB+
Perfhi
IFNγ+
I.
% Hepatic Dendritic cells
Day 8
Day 31
% of CD45+ CD4- cells
WT
CD4 deplete
cDC1
cDC2
J.
### Hepatic Dendritic cells
Day 8
Day 31
### cDCs
WT
CD4 deplete
cDC1
cDC2
K.
MHCII Expression on cDCs
Day 8
Day 31
MHCII MFI
WT
CD4 deplete
cDC1
cDC2
L.
Adult (WT splenocytes)
100 µm
M.
Adult (CD4-deplete splenocytes)
100 µm
N.
Young (WT splenocytes)
100 µm

**Fig. S5: Young mice and CD4-depleted mice exhibit impaired CD8+ T cell function accompanied by a decrease in cDC activity.** (**A-F**) Hepatic leukocytes from adult (black circles) and young (blue triangles) HBVtgRag-/- mice adoptively transferred with WT splenocytes were assessed by flow cytometry at the indicated timepoints. (**A**) Number of $sAg_{353}$+ (solid lines) and $sAg_{370}$+ (dashed lines) CD8+ T cells. (**B**) Percentage of effector/effector memory $sAg_{370}$+ T cells; CD44+ CD62L-. (**C**) Number of GzmB+, Perf+, and IFNγ+ $sAg_{353}$+ CD8+ T cells. (**D**) Percentage of total (filled bars & symbols) and $sAg_{370}$+ (open bars & symbols) CD8+ T cells expressing markers associated with T cell exhaustion: TOX, PD-1, CD27, CD39, TIGIT, TIM3, LAG-3, VISTA. Percentage of total and $sAg_{370}$+ CD8+ T cells expressing (**E**) effector molecules GzmB and perforin and (**F**) memory/proliferative capacity markers TCF-1 and CD127. (**G-N**) Adult mice were adoptively transferred with WT splenocytes (WT; black circles) or CD4-deplete WT splenocytes (CD4-deplete; pink squares) and hepatic leukocytes were assessed by flow cytometry on day 8 and 31 or by imaging on day 8. (**G**) Number of $sAg_{353}$+ and $sAg_{370}$+ CD8+ T cells. (**H**) Number of total, $sAg_{353}$+, and $sAg_{370}$+ CD8+ T cells that were GzmB+, Perf+ or IFNγ+. (**I**) Percentage and (**J**) number of cDC1 and cDC2. (**K**) MHCII MFI on cDC1 and cDC2. CD4 (yellow) and CD8 (purple) staining of adult (**L**), CD4-deplete (**M**), and young (**N**) livers on day 8. (A-B) N≥9 mice combined from three independent experiments, (C) N≥6 mice, representative of two independent experiments, (D-N) N≥4, representative of two independent experiments; statistics performed using unpaired t-test without multiple test correction.

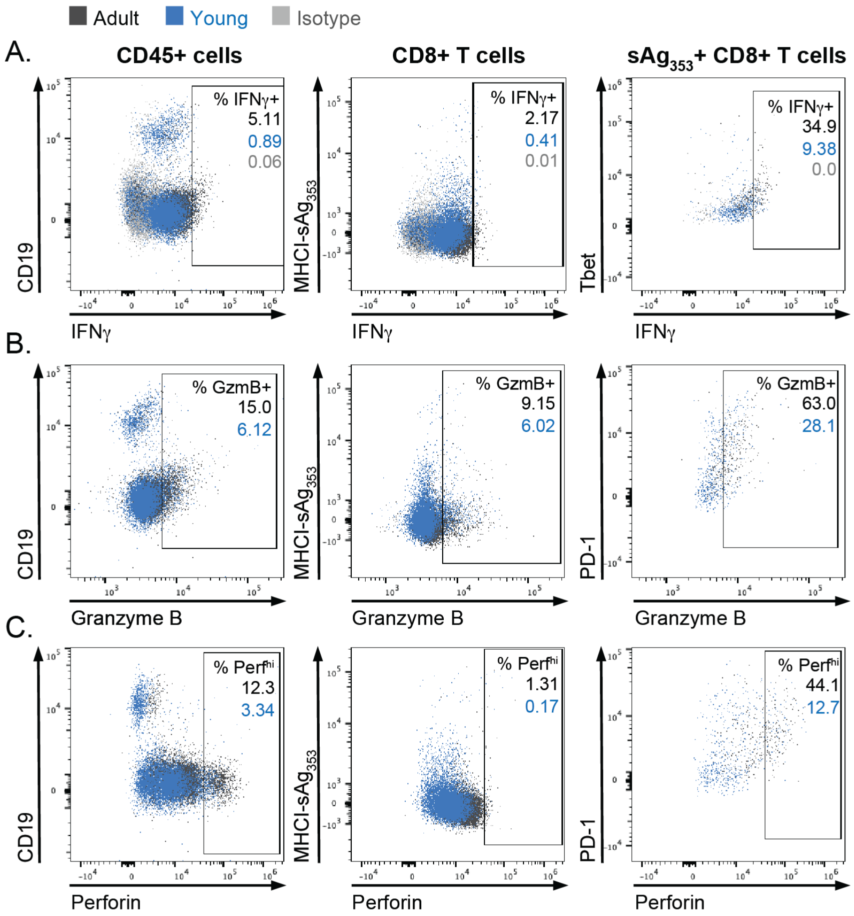


**Fig. S6: Representative gating demonstrating that CD8+ T cells in adult mice express higher levels of IFNγ, Granzyme B, and Perforin compared to CD8+ T cells in young mice.** Hepatic leukocytes from adult (black) and young (blue) HBVtgRag-/- mice adoptively transferred with adult WT splenocytes were assessed by flow cytometry and compared to isotype-stained controls (gray). Representative intracellular cytokine staining of (**A**) IFNγ, (**B**) Granzyme B, and (**C**) Perforin on (left) total lymphocytes relative to CD19, (middle) CD8+ T cells relative to MHCI-sAg$_{353}$, and (right) sAg$_{353}$+ CD8+ T cells relative to PD-1.

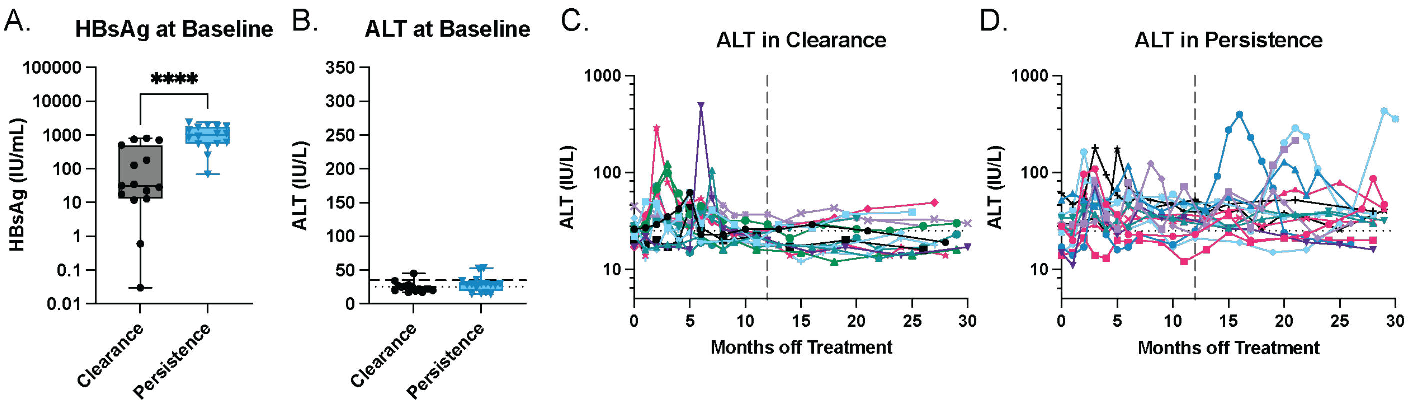


**Fig. S7: Baseline HBsAg levels and dynamics of ALT elevation in study patients.** (**A**) Quantitative HBsAg and (**B**) ALT levels at baseline in HBsAg clearance (black circles) vs. persistence (blue triangles); and over time in the (**C**) HBsAg clearance and (**D**) HBsAg persistence groups, shown with individual patients in different colors and symbols. N ≥ 15 per group; statistics performed with unpaired t-test.

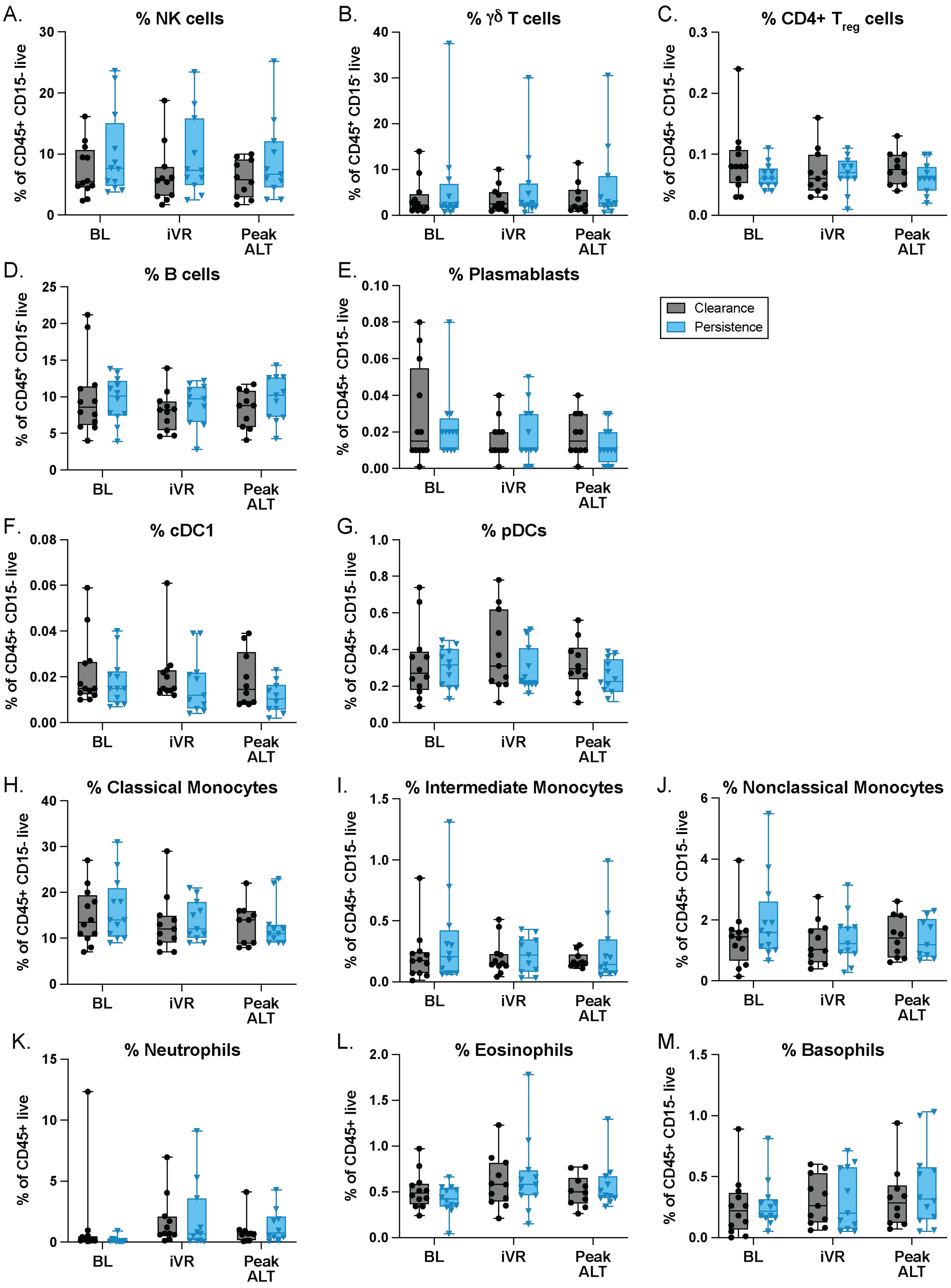

A.
% NK cells
% of CD45+ CD15- live
30
20
10
0
BL
iVR
Peak ALT
B.
% γδ T cells
% of CD45+ CD15- live
40
30
20
10
0
BL
iVR
Peak ALT
C.
% CD4+ Treg cells
% of CD45+ CD15- live
0.3
0.2
0.1
0.0
BL
iVR
Peak ALT
D.
% B cells
% of CD45+ CD15- live
25
20
15
10
5
0
BL
iVR
Peak ALT
E.
% Plasmablasts
% of CD45+ CD15- live
0.10
0.08
0.06
0.04
0.02
0.00
BL
iVR
Peak ALT
Clearance
Persistence
F.
% cDC1
% of CD45+ CD15- live
0.08
0.06
0.04
0.02
0.00
BL
iVR
Peak ALT
G.
% pDCs
% of CD45+ CD15- live
1.0
0.8
0.6
0.4
0.2
0.0
BL
iVR
Peak ALT
H.
% Classical Monocytes
% of CD45+ CD15- live
40
30
20
10
0
BL
iVR
Peak ALT
I.
% Intermediate Monocytes
% of CD45+ CD15- live
1.5
1.0
0.5
0.0
BL
iVR
Peak ALT
J.
% Nonclassical Monocytes
% of CD45+ CD15- live
6
4
2
0
BL
iVR
Peak ALT
K.
% Neutrophils
% of CD45+ live
15
10
5
0
BL
iVR
Peak ALT
L.
% Eosinophils
% of CD45+ live
2.0
1.5
1.0
0.5
0.0
BL
iVR
Peak ALT
M.
% Basophils
% of CD45+ CD15- live
1.5
1.0
0.5
0.0
BL
iVR
Peak ALT

**Fig. S8: Frequencies of immune cell subsets that are similar in HBsAg clearance and persistence patients.** PBMCs were stained and assessed by CyTOF to measure frequency of the indicated immune cell subsets at respective timepoints in cases with HBsAg clearance (black circles) and HBsAg persistence (blue triangles). Percentage of (**A**) NK cells, (**B**) γδ T cells, (**C**) CD4+ Treg cells, (**D**) B cells, (**E**) Plasmablasts, (**F**) cDC1, (**G**) pDCs, (**H**) Classical Monocytes, (**I**) Intermediate Monocytes, (**J**) Non-classical Monocytes, (**K**) Neutrophils, (**L**) Basophils, and (**M**) Eosinophils in total CD15- PBMCs. $N \geq 9$ per group per timepoint; statistics performed using multiple unpaired t-tests with Benjamini Hochberg FDR correction ($Q = 0.1$; $q > Q$ for all comparisons).

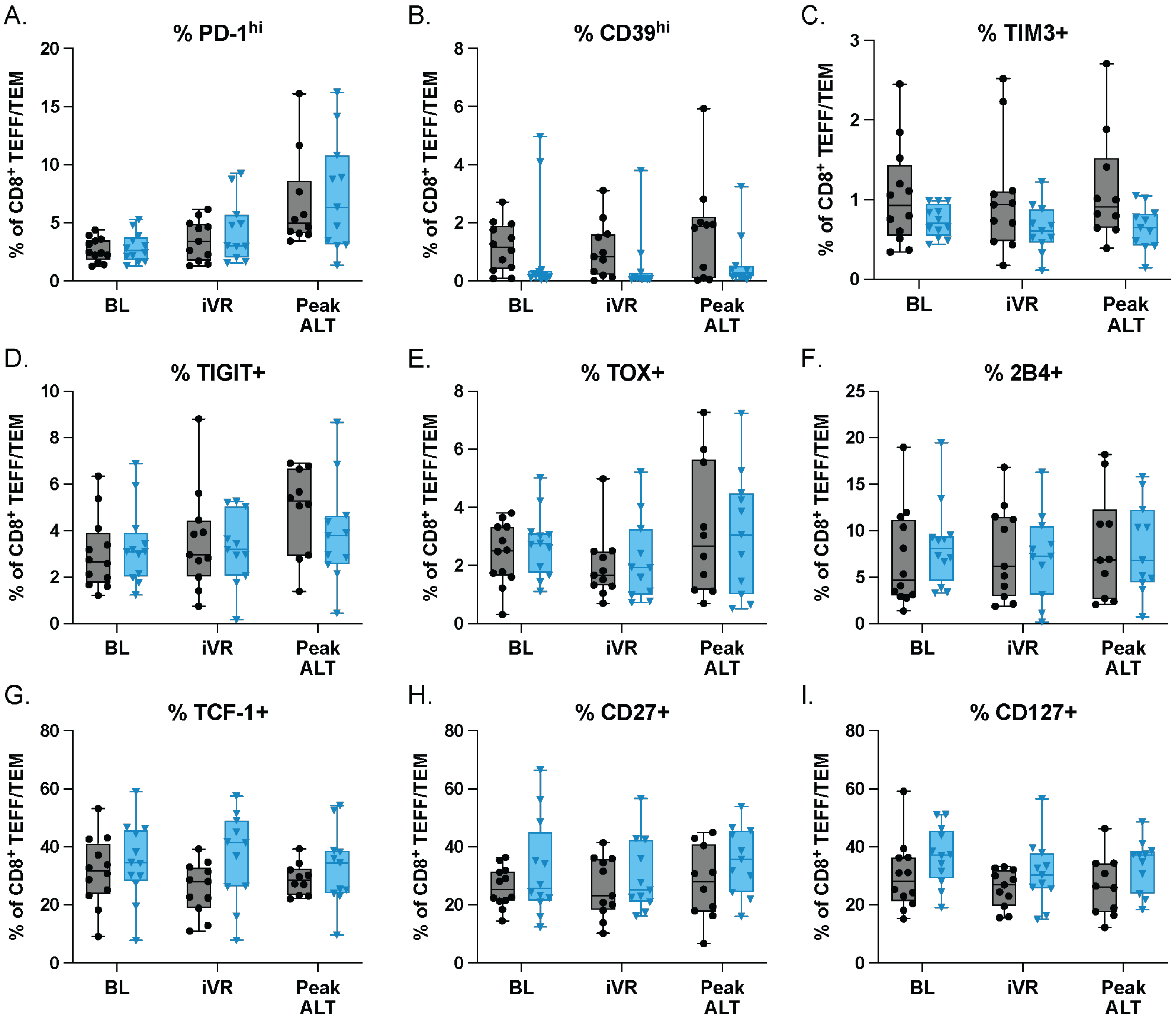


**Fig. S9: Peripheral CD8+ T cells from CHB patients that clear or retain circulating HBsAg do not differ in their expression of canonical "exhaustion/dysfunction" markers.** (**A-I**) PBMCs from patients with HBsAg clearance (black circles) and persistence (blue triangles) after stopping NA were stained at the indicated timepoints and assessed by CyTOF to measure the expression of markers associated with T cell exhaustion/dysfunction as well as markers of stemness and memory on effector/memory ($T_{EFF/EM}$, all except CD45RA+ CCR7+) CD8+ T cells. Percentage of CD8+ $T_{EFF/EM}$ cells that are (**A**) PD-1$^{hi}$, (**B**) CD39$^{hi}$, (**C**) TIM3+, (**D**) TIGIT+, (**E**) TOX+, (**F**) 2B4+, (**G**) TCF-1+, (**H**) CD27+, and (**I**) CD127+ at the indicated timepoints. N≥9 group per timepoint; statistics performed using multiple unpaired t-tests with Benjamini Hochberg FDR correction ($Q = 0.1$; $q > Q$ for all comparisons).

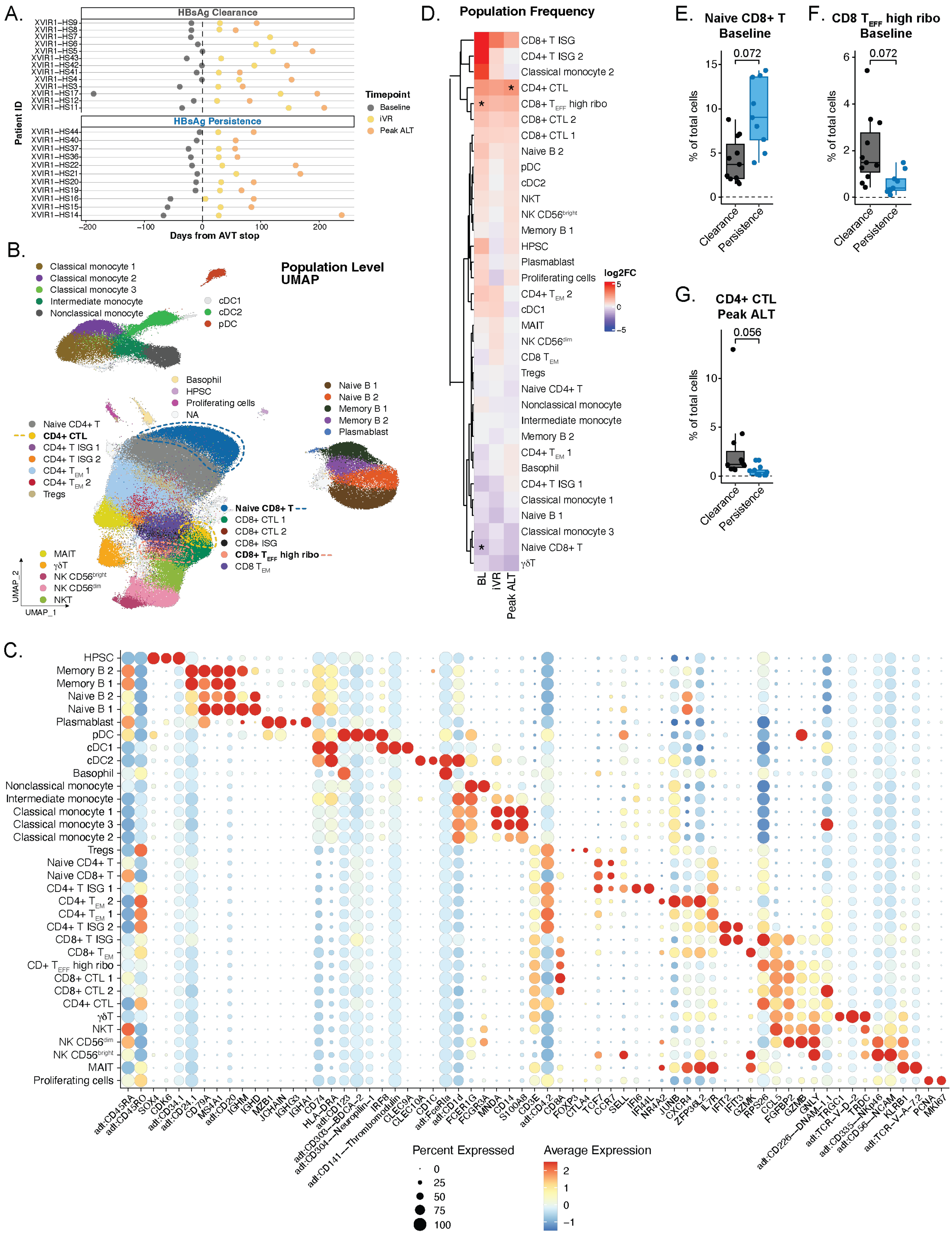
A.
HBsAg Clearance
XVIR1–HS9
XVIR1–HS8
XVIR1–HS7
XVIR1–HS6
XVIR1–HS5
XVIR1–HS43
XVIR1–HS42
XVIR1–HS41
XVIR1–HS4
XVIR1–HS3
XVIR1–HS17
XVIR1–HS12
XVIR1–HS11
HBsAg Persistence
XVIR1–HS44
XVIR1–HS40
XVIR1–HS37
XVIR1–HS36
XVIR1–HS22
XVIR1–HS21
XVIR1–HS20
XVIR1–HS19
XVIR1–HS16
XVIR1–HS15
XVIR1–HS14
Patient ID
Days from AVT stop
−200
−100
0
100
200
Timepoint
Baseline
iVR
Peak ALT
B.
Population Level UMAP
Classical monocyte 1
Classical monocyte 2
Classical monocyte 3
Intermediate monocyte
Nonclassical monocyte
cDC1
cDC2
pDC
Basophil
HPSC
Proliferating cells
NA
Naive B 1
Naive B 2
Memory B 1
Memory B 2
Plasmablast
Naive CD4+ T
CD4+ CTL
CD4+ T ISG 1
CD4+ T ISG 2
CD4+ T EM 1
CD4+ T EM 2
Tregs
Naive CD8+ T
CD8+ CTL 1
CD8+ CTL 2
CD8+ ISG
CD8+ T EFF high ribo
CD8 T EM
MAIT
γδT
NK CD56bright
NK CD56dim
NKT
UMAP_2
UMAP_1
C.
HPSC
Memory B 2
Memory B 1
Naive B 2
Naive B 1
Plasmablast
pDC
cDC1
cDC2
Basophil
Nonclassical monocyte
Intermediate monocyte
Classical monocyte 1
Classical monocyte 3
Classical monocyte 2
Tregs
Naive CD4+ T
Naive CD8+ T
CD4+ T ISG 1
CD4+ T EM 2
CD4+ T EM 1
CD4+ T ISG 2
CD8+ T ISG
CD8+ T EM
CD+ T EFF high ribo
CD8+ CTL 1
CD8+ CTL 2
CD4+ CTL
γδT
NKT
NK CD56dim
NK CD56bright
MAIT
Proliferating cells
adt:CD45RA
adt:CD45RO
SOX4
CDK6
adt:CD34.1
adt:CD24.1
CD79A
MS4A1
adt:CD20
IGHM
IGHD
MZB1
JCHAIN
IGHG3
IGHA1
CD74
HLA–DRA
adt:CD123
adt:CD303—BDCA–2
adt:CD304—Neuropilin–1
adt:CD141—Thrombomodulin
IRF8
CLEC9A
CLEC10A
CD1C
adt:FceRIa
adt:CD1d
FCER1G
FCGR3A
MNDA
CD14
S100A8
CD3E
adt:CD4.2
CD8A
FOXP3
CTLA4
TCF7
CCR7
SELL
IFI6
IFI44L
NR4A2
JUNB
CXCR4
ZFP36L2
IL7R
IFIT2
IFIT3
GZMK
RPS26
CCL5
FGFBP2
GZMB
GNLY
adt:CD226—DNAM–1.1
TRGC1
adt:TCR–V–D–2
TRDC
adt:CD335—NKp46
adt:CD56—NCAM
KLRB1
adt:TCR–V–A–7.2
PCNA
MKI67
Percent Expressed
0
25
50
75
100
Average Expression
2
1
0
−1
D.
Population Frequency
CD8+ T ISG
CD4+ T ISG 2
Classical monocyte 2
CD4+ CTL
CD8+ T EFF high ribo
CD8+ CTL 2
CD8+ CTL 1
Naive B 2
pDC
cDC2
NKT
NK CD56bright
Memory B 1
HPSC
Plasmablast
Proliferating cells
CD4+ T EM 2
cDC1
MAIT
NK CD56dim
CD8 T EM
Tregs
Naive CD4+ T
Nonclassical monocyte
Intermediate monocyte
Memory B 2
CD4+ T EM 1
Basophil
CD4+ T ISG 1
Classical monocyte 1
Naive B 1
Classical monocyte 3
Naive CD8+ T
γδT
BL
iVR
Peak ALT
log2FC
5
0
−5
E.
Naive CD8+ T
Baseline
0.072
% of total cells
15
10
5
0
Clearance
Persistence
F.
CD8 T EFF high ribo
Baseline
0.072
6
4
2
0
Clearance
Persistence
G.
CD4+ CTL
Peak ALT
0.056
% of total cells
10
5
0
Clearance
Persistence

**Fig. S10: CITE-seq sample collection, single cell population identity, and population frequency.** Single-cell sequencing using CITE-seq was performed on PBMCs from 24 patients (n=13 HBsAg clearance, n=11 HBsAg persistence) to detect mRNA transcripts and selected proteins. (**A**) Depiction of clinical sampling times relative to treatment withdrawal in HBsAg clearance and persistence patient groups. (**B**) Uniform manifold approximation and projection (UMAP) visualization individual cells clustered by a weighted nearest neighbor (WNN) approach using a combination of RNA and protein reads at the population level. (**C**) Heatmap depicting the expression of various genes and ADTs (protein-level expression) in each cluster compared to all other clusters determined by MAST and useful for cluster annotation. Dot color represents gene/ADT expression level and dot size represents the percentage of cells in a given cluster expressing the gene/ADT. (**D**) Heatmap showing $\log_2$fold-difference enrichment of individual clusters at the population level in Clearance (red) versus Persistence (blue) BeNEG-DO patients at each indicated timepoint. Populations with a significantly different frequency (adj.p<0.1) are marked with *. Boxplots depicting the frequencies of (**E**) Naïve CD8+ T cells at baseline, (**F**) CD8+ $T_{EFF}$ cells at baseline, and (**G**) CD4+ CTL at peak ALT, the only three populations with significant differences in their frequency between clearance (black) versus persistence (blue) patients.

Table S1. Patient Characteristics

| | **Total n=31** | **HBsAg Clearance n=15** | **HBsAg Persistence n=16** | **P value** |
|---|---|---|---|---|
| Female, n (%) | 11 (35.48) | 2 (13.3) | 9 (56.25) | 0.02 |
| #Age, years | | | | 0.33 |
| Median (25th-75th%-ile) | 51 (47.5-59) | 51 (48.5-62.5) | 50 (46.2-58.2) | |
| Range | 31-67 | 47-66 | 31-67 | |
| Race, n (%) | | | | >0.999 |
| Asian | 31 (100.0) | 15 (100.0) | 16 (100.0) | |
| Continent born, n (%) | | | | 0.6 |
| Asia | 28 (90.3) | 13 (86.7) | 15 (93.75) | |
| North America | 3 (9.7) | 2 (13.3) | 1 (6.25) | |
| Presumed mode of HBV transmission, n (%) | | | | 0.2 |
| Vertical | 20 (64.5) | 8 (53.3) | 12 (75.0) | |
| Unknown | 11 (35.5) | 7 (46.7) | 4 (25.0) | |
| HBV genotype | | | | >0.999 |
| A | 2 (6.5) | 1 (6.7) | 1 (6.25) | |
| B | 19 (61.3) | 9 (60.0) | 10 (62.5) | |
| C | 10 (32.2) | 5 (33.3) | 5 (31.25) | |
| *Circulating HBsAg (IU/mL) at BL Median (25th-75th%-ile) | 541.84 (33-997.4) | 31.5 (15.2-338.3) | 997.4 (557.2-1721.9) | <0.001 |
| Range | 0.03-2419.6 | 0.03-803.4 | 69-2419.6 | |

All patients were HBsAg positive/anti-HBs negative and HBeAg negative/anti-HBe positive, with serum HBV DNA <20 IU/mL when starting and stopping NA treatment. NA therapy included: Entecavir (n=12), Tenofovir (n=10), Tenofovir and Entecavir (n=3). All patients with bridging liver fibrosis (>F2) were excluded from study participation.

#Age at the time of treatment withdrawal.

*qHBsAg assay (Abbott Diagnostics Division, Sligo, Ireland).

Fisher's exact test was used for categorical data and two-tailed unpaired Student's *t* test for continuous data. $P < 0.05$ was considered significant.

**Supplemental Materials & Methods:**

**Mice and experimental system**

All mice were bred and housed in specific pathogen–free housing under an IACUC-approved protocol (AN170936) and in accordance with the guidelines of the Laboratory Animal Resource Center of the University of California, San Francisco. WT C57BL/6 mice were purchased from Jackson Laboratory and subsequently bred at UCSF. HBVtg*Rag1*$^{-/-}$ mice (including HBVEnv*Rag1*$^{-/-}$ and HBVRpl*Rag1*$^{-/-}$ strains) were previously described (*14*). Briefly, HBVEnv*Rag1*$^{-/-}$ mice were generated using HBV-Envelope mice [lineage 107-5D; gift from F. Chisari, Scripps Research Institute (*45*)] backcrossed to *Rag1*$^{-/-}$ C57BL/6 mice for 15 generations. HBVEnv*Rag1*$^{-/-}$ mice contain the entire envelope (subtype ayw) protein-coding region under the constitutive transcriptional control of the mouse albumin promoter. HBVRpl*Rag1*$^{-/-}$ mice were generated using HBV replication mice [lineage 1.3.46; gift from F. Chisari (*46*)] crossed to *Rag1*$^{-/-}$ C57BL/6 mice for 15 generations. HBVRpl*Rag1*$^{-/-}$ mice contain a terminally redundant HBV DNA construct and produce infectious genotype D human HBV in hepatocytes and in the proximal convoluted tubules of their kidneys. Young (3-3.5 weeks old, pre-weaning) or adult (>8 weeks old) HBVtg*Rag1*$^{-/-}$ mice were given 0.8-1.0x$10^8$ syngeneic splenocytes pooled from adult (>8 weeks old) WT mice in 0.5 ml of saline via tail vein injection. Mice were bled at regular intervals on days 5, 7, 14, 21, 28, 56, and 84, unless otherwise specified. Mice were followed for plasma ALT using an ALT-SL kit (Sekisui Diagnostics) on a Cobas Mira Plus analyzer (Roche Diagnostics).

**HBV protein assays**

Plasma was collected and assayed for the presence of HBsAg using the Hepatitis B Surface Antigen ELISA kit (Creative Diagnostics) and measured on an ELx800 plate reader (BioTek). HBsAg results are reported as positive or negative following manufacturer instructions. Total anti-HBs was quantified using the Antibody to HBsAg ELISA kit with standards (Creative Diagnostics).

**Cell preparations**

Leukocytes were isolated from the liver after perfusion and digestion. For experiments targeting myeloid cells, mice were perfused via the inferior vena cava with 30 mL of digestion media [HBSS with collagenase type IV (92.53 units/mL; Worthington Biochemical Corporation), and DNase I (0.02 mg/ml; Roche Diagnostics)]. Livers were chopped and further digested with liberase and DNase I (1 Wünsch Unit (WU) and 0.8 mg, respectively] for 30 min at 37°C while shaking at 225 rpm. For experiments targeting lymphocyte populations, mice were perfused to remove blood within the liver (<1 min); no additional enzymatic digestion was performed. For both preparations, livers were forced through a 70 μm cell strainer for lymphocyte preparation and 100 μm cell strainer for myeloid preparations, and debris was removed by centrifugation at 30 *g* for 3 min. Supernatants were collected and centrifuged for 10 min at 650 *g*. Cells were resuspended in RPMI 1640 with glutamine, 5% FBS, and 40% Percoll® (GE Healthcare) and then underlaid with 60% Percoll®, followed by centrifugation for 20 min at 1450 *g*. Leukocytes were isolated from the Percoll® interface, washed and counted on a Luna-FL™ dual fluorescence cell counter (Logos Biosystems) using acridine orange/propidium iodide (Logos Biosystems).

Splenocytes were isolated by grinding tissue between two frosted microscope slides and filtered through a 70 μm cell strainer with cold PBS. For flow cytometry experiments, cell suspensions were pelleted and lysed with ACK lysing buffer for 3-5 min to remove red blood cells and subsequently washed into staining buffer and counted. For adoptive transfer cell preparations, splenocytes were not lysed and instead filtered and washed into sterile saline at a concentration of 1.6 – 2x$10^8$ cells/mL.

#### Antibodies and Flow Cytometry

Hepatic leukocytes and splenocytes were prepared as above. One million cells were stained according to standard protocols in MACS staining buffer (PBS with 0.5% BSA and 2 mM EDTA) with combinations of anti-mouse antibodies listed below. Briefly, cells were stained with viability dye (Live/Dead Aqua, Zombie Aqua, or Zombie NIR) for 20 min at room temperature, Fc-blocked with anti-CD16/-CD32 for 10 min at 4°C and stained for 15 min at 4°C with a surface antibody cocktail.

For experiments with intracellular cytokine detection, cells were treated with Brefeldin A (1 μg/mL) at 37°C and 5% $CO_2$ for 4 hours prior to surface staining and fixation. Cells were fixed and permeabilized using the eBioscience™ FoxP3/Transcription Factor Staining Buffer kit and stained overnight at 4°C with intracellular antibodies targeting cytokines and/or transcription factors. In some experiments, IFNγ secretion assays (Miltenyi Biotech) were used to detect cells actively secreting IFNγ using a capture antibody with secondary detection system directly *ex vivo* without stimulation.

For experiments using MHCII tetramers, prior to the viability stain, cells were washed twice with PBS and stained for 1 hour at 37°C and 5% $CO_2$ with 0.5 ug of MHCII-tetramers. For experiments using MHCI tetramers, prior to the viability stain (and after the MHCII stain, if applicable), cells are stained for 1hr at 4°C in staining buffer with 0.7 ug (MHCI-$sAg_{353}$) or 1.5 ug (MHCI-$sAg_{370}$) of MHCI tetramer. The threshold of detection for the purposes of defining a sAg-specific response was considered as twice the frequency of tetramer$^+$ T cells seen in HBV- (no transgene) mice.

Cells were analyzed immediately after completing staining using either a LSR II flow cytometer (BD Biosciences), Aurora spectral flow cytometer (Cytek), or, for cell sorting experiments, an Aria III (BD Biosciences). For BD Bioscience instruments, data was collected using FACSDiva software and analyzed using FlowJo (BD Biosciences). For the Cytek spectral flow cytometer, data was collected and preprocessed to correct spillover matrices using SpectroFlo® software with subsequent analysis using FlowJo.

#### Chromogenic immunohistochemistry (IHC) tissue staining and imaging

Liver tissue was fixed in formalin for 24 hours (2hr at 4°C then 22hr at room temp.) and embedded in paraffin blocks at the UCSF Biorepository and Tissue Biomarker Technology Core. Three 4 μm serial sections per sample were cut and mounted on a slide.

Tissue sections were stained with rabbit anti-mouse CD4 (EPR19514, Abcam, 1:1000), rat anti-mouse CD8α (4SM15, eBioscience/Invitrogen, 1:50), and rabbit anti-mouse RORγ (EPR20006, Abcam, 1:3000) using the Ventana Discovery Ultra staining platform. Antigen retrieval and tissue conditioning were completed using CC1 conditioning at 97°C for 32 min with 16 min goat Ig blocking followed by CC2 stripping at 97°C for 8 min. Anti-RORg was stained first at 36°C for 32 min + 24 min with OmniMap anti-Rabbit secondary, then purple chromogen detection for 40 min, followed by anti-CD8a at 36°C for 32 min + 24 min OmniMap anti-Rat secondary, then teal chromogen detection for 32 min, and lastly by anti-CD4 at 36°C for 32 min + 12 min OmniMap anti-Rabbit-NP secondary + 12 min secondary enzyme conjugate and yellow chromogen detection for 20 min. Sections were dried and coverslipped followed by imaging at 20x resolution on the Zeiss Axioscanner Z1. Native, uncompressed CZI imaging files were used as a starting point for subsequent image analysis.

#### IHC imaging analysis

CZI images were converted to the Big Tiff file format using Zen software (Zeiss), and then to HDF5 format using the Ilastik plug-in for FIJI (*47-49*). We trained two Random Forest classifiers in the 1.3.3 version of Ilastik (*50*). The training set comprised 58 2000x2000-pixel images taken across four tissue samples from the RORγ/CD8α/CD4 triplex stain. One classifier is a binary classifier used to predict nuclei pixels and stroma pixels in an RGB image. The nuclei pixels were segmented into objects using hysteresis thresholding (Gaussian smoothing σ = 1.0, core threshold = 0.55, final threshold = 0.65). The second classifier was trained to classify these objects into one of 6 cell types: CD4, CD8, Th17 (CD4+ RORγ+), ILC3 (RORγ+), Tc17 (CD8+ RORγ+) and "other". The output of this classifier provided the predicted label and the coordinates of the center of these objects.

We built a GUI to take these coordinates and labels as input, so that they could be visualized and filtered based on user-provided parameters. The GUI also allows users to create custom regions of interest (ROIs) on the raw image to filter out tissue and staining artefacts. Using the GUI, we filtered the output of each sample prediction to include only immune cells (i.e. removed "other" cell types) and remove tissue artefacts. We used the scikit-learn (*51*) Python implementation of Density-Based Spatial Clustering of Applications with Noise (DBSCAN) (*52*) to cluster the filtered immune cell objects. All parameters used were defaults, except for the "eps" parameter, which is "the maximum distance between two samples for one to be considered as in the neighborhood of the other," which we specified as 100 pixels; clusters were defined as ≥5 cells within a neighborhood.

**_In vitro_ T cell priming assays**

Antigen presenting cells were purified using fluorescence activated cell sorting (FACS) of hepatic or splenic cell suspensions pooled from the livers of ≥ 8 adult or young resting, naïve mice. Separately, naïve OT-II $CD4^+$ T cells were FACS-enriched from the spleens of adult mice. $5x10^3$ sorted APCs were plated in a 96-well round-bottom plate and loaded with 10 μg/mL ovalbumin peptide ($Ova_{323-339}$) in culture media (RPMI with glutamine + 10% FBS + 1x β-mercaptoethanol) for 1 hour at 37°C and 5% $CO_2$. Naïve OT-II cells were labeled with 1 μM CFSE for 5 min at 37°C. $5x10^3$ Ova-loaded APCs were co-cultured with $1.5x10^4$ naïve OT-II $CD4^+$ T cells for 60 hr. Supernatants were stored at -80°C for analysis of cytokine secretion using Cytometric Bead Array Flex Kits (BD Biosciences). Before recovering co-cultured cells, $10^6$ *Rag*$^{-/-}$ splenocytes were spiked into each well to enable reproducible recovery, staining, and analysis.

**CD4 depletion**

Wild-type adult (>8 wks) splenocytes were prepared as described above and split into two with one portion set-aside at 4°C untouched until ready for injection while the other portion was depleted of CD4+ cells using anti-CD4 (L3T4) microbeads with LD MACS columns (Miltenyi Biotec). The same number of sex-matched wild-type or CD4-depleted splenocytes ($1.0*10^8$ cells) were then adoptively transferred into HBVtgRag-/- recipient adult (>8 wks) mice. Depletion efficiency (> 99%) was checked by flow cytometry by staining an aliquot of donor splenocytes prior to transfer.

**Human Subjects Sample collection and PBMC isolation**

Longitudinal blood samples were obtained under the institutional review board (IRB) protocol #11-07994 at UCSF and Sutter Health for the BeNEG-DO study and protocol 2012.059-2 (SCoo) at Sutter Health for the BeNEG pilot patient. Synchronous clinical and research blood samples were collected monthly for the first 6 months after NA withdrawal every 2 months during months 7-12, and every 3 months beyond month 12. Lab monitoring frequency exceeded protocol during ALT flares (ALT≥200 U/L) and/or at the discretion of the managing hepatologist. HBsAg seroclearance was defined when qHBsAg <.05 IU/mL and was undetected by standard qualitative assay conducted by the clinical laboratory. HBsAg- status was confirmed, initially within 3-months, then every 6 months as applicable.

A healthy donor sample purchased from Vitalant Research Institute (San Francisco, CA) was used as a batch control for CyTOF experiments. Blood was collected into sterile BD Vacutainer® EDTA Tubes (BD Biosciences) and centrally processed and stored in the Ibrahim El-Hefni Liver Biorepository. Peripheral Blood Mononuclear Cells (PBMC) were isolated from whole blood using Ficoll-Paque PLUS (GE Healthcare Pharmacia) density gradient and centrifuged at 400 *g* for 30 min without braking. PBMC were collected from the Ficoll-Paque PLUS interface, washed twice with Dulbecco's PBS supplemented with 2mM EDTA, and aliquots of $5x10^6$ PBMC in 0.5 mL of freezing medium (9:1 FBS/DMSO) were stored in liquid nitrogen.

**HBV genotyping**

Full length hepatitis B virus (HBV) sequences were derived from BeNEG-DO patient plasma samples. HBV DNA was isolated from serum obtained during viremia (QIAmp DNA blood mini kit, Qiagen; threshold limit of sequence detection is ~500 viral DNA copies/mL). Amplification of HBV DNA was performed by nested PCR adapted from a previously described method (*53*). The PCR product was purified (Qiagen QIAquick PCR purification kits) and the purified amplicons were sequenced using the following primers: (1a) 251F, 1190R (940bp), (1b) 595F, 1190F (1203bp), (2a) 2300F, 215R (1131bp), (2b) 2819F, 617R (1032bp), (3) 1877F, 2835R (959bp) and (4) 2331R (748bp) (Elim Biopharmaceuticals, Hayward, CA). Full length viral sequences were assembled and aligned using Lasergene 15.0 (DNASTAR). The sequences were annotated and genotyped using The Hepatitis B Virus Database (*54*).

**Mass cytometry (CyTOF) data generation and analysis**

A 41-parameter CyTOF panel was designed to assess major immune populations in PBMC and markers of T cell activation/inhibition/exhaustion. Mass cytometry was performed as described (*55*). Metals were conjugated using the MaxPAR antibody conjugation kit (Fluidigm) according to the manufacturer's instructions. Each antibody clone was titrated to optimal staining concentrations using unstimulated or anti-CD3/CD28 stimulated healthy donor PBMC.

Thawed PBMC samples with viability >75% determined using a Vi-Cell XR cell counter (Beckman Coulter) were used for cell staining. Briefly, $2.5x10^6$ cells were incubated with 50µM Cisplatin (Sigma-Aldrich, San Luis, MO) for 60s before quenching 1:1 with PBS with 0.5% BSA and 5mM EDTA to label dead cells. Cells were centrifuged at 500 *g* and resuspended in cell staining media (CSM) (Fluidgim) and blocked with Human TruStain FcX™ block (BioLegend) for 5 min at room temperature, followed by staining with anti-CXCR5 antibody in CSM (3mg/mL) for 30 min at 4°C while shaking at 90rpm. Cells were fixed using the Cell-ID™ 20-Plex Pd Barcoding Kit (Fluidigm) for 10 min at room temperature and barcoded by mass-tag labeling with distinct combinations of stable Pd isotopes as previously described (*56*). Twenty barcoded samples were pooled together for staining with surface markers antibodies for 30 min at room temperature while shaking. Cells were permeabilized with Permeabilization buffer (eBioscience) followed by intracellular staining for one hour at 4°C while shaking. Finally, cells were stained with 191/193Ir DNA intercalator (Fluidigm) diluted in 1.6% PFA at 4 °C overnight. Cells were washed and resuspended at $1.2x10^6$ cells/mL in deionized water + 10% EQ four element calibration beads (Fluidigm) and run on a Fluidigm CyTOF2 Helios mass Cytometer within one week of staining.

CyTOF fcs files were normalized to the EQ™ calibration beads, concatenated, and de-barcoded using premessa pipeline (https://github.com/ParkerICI/premessa) (*57*). CellEngine software (CellCarta) was used to exclude the EQ™ calibration beads and manually gate the files. Singlets were gated by DNA signal (Ir191) and Event Length. Live cells were identified as cisplatin Pt195 negative. CD45- Pt195+ cells were excluded from the analysis. Potential batch effects were minimized by including a control sample from the same individual in each of 9 experimental runs.

**Bulk RNA-seq**

RNA was extracted from 1-2.5x$10^6$ PBMCs using Qiagen RNeasy mini kits (Qiagen), and integrity was measured with the Agilent 2100 Bioanalyzer (Agilent Technologies). Ribosomal and hemoglobin depleted total RNA-sequencing libraries were created using Universal plus mRNA with NuQuant® (TECAN), with changes to the protocol as follows: First, the Poly(A) Selection step was omitted. Second, RNA Fragmentation and QIAseq FastSelect (Qiagen) steps were combined to remove rRNA and globin from the RNA samples. Third, 10ul of total RNA was combined with 1X TECAN Fragmentation Buffer. The rest of the Universal plus mRNA with NuQuant® protocol was followed according to vendor instructions starting with First strand cDNA synthesis step and skipping the optional steps of AnyDeplete and NuQuant with automation on a Beckmen BioMek FXp system. Libraries were subsequently normalized and pooled using a Labcyte Echo 525 system (Beckman Coulter). The pooled libraries were sequenced on an Illumina NovaSeq 6000 S4 flow cell lane with paired end 150 bp reads (~29.5M reads/sample).

**Computational processing of Bulk RNA-Seq data for genotyping**

Sequencing reads were aligned to the human reference genome and Ensembl annotation (GRCh38 genome build, version 95) using STAR (v2.7.5c) (*58*) with the following parameters: --outFilterType BySJout --outFilterMismatchNoverLmax 0.04 -outFilterMismatchNmax 999 --alignSJDBoverhangMin 1 --outFilterMultimapNmax 1 -alignIntronMin 20 --alignIntronMax 1000000 --alignMatesGapMax 1000000. Duplicate reads were removed using Picard Tools (v2.23.3) (*59*). Nucleotide variants were identified from the resulting bam files using the Genome Analysis Tool Kit (v4.0.11.0) following the best practices for RNA-seq variant calling (*60, 61*). This included splitting spliced reads, calling variants with HaplotypeCaller (added parameters: --dont-use-soft-clipped-bases -standcall-conf 20.0), and filtering variants with VariantFiltration (added parameters: -window 35 cluster 3 --filter-name FS -filter FS > 30.0 --filter-name QD -filter QD < 2.0). Variants were further filtered to include a list of high quality SNP for identification of the subject of origin of individual cells by removing all novel variants, maintaining only biallelic variants with MAF greater than 5%, a max missing of one individual with a missing variant call at a specific site and requiring a minimum depth of two (parameters: --max-missing 1.0 --min-alleles 2 --max-alleles 2 --removeindels --snps snp.list.txt --min-meanDP 2 --maf 0.05 --recode --recode-INFO-all –out).

**CITE-seq sample loading and sequencing**

A total of 153 samples from 24 patients representing longitudinal samples for each individual were multiplexed for CITE-seq. PBMC suspensions from 16 individual patients were pooled together with equivalent numbers of live cells, determined using Vi-CELL XR (Beckman Coulter) or Cellaca MX (Nexcelom Bioscience) cell counters. For samples with viability <55% (6% of samples) live cells were enriched with the Dead Cell Removal kit (Miltenyi Biotec). 1x$10^6$ pooled cells were Fc blocked (Human TruStain FcX, BioLegend) for 10 min at room temperature followed by staining with a custom TotalSeq™-A antibody panel (PG00022; BioLegend; data file S3) for 45 minutes at 4°C. Cells were washed 3 times with Cell Staining Buffer (BioLegend) and resuspended in DPBS (w/o $Ca2^+$, $Mg2^+$) with 0.04% BSA at 2,000 cells/µL. 65,000 pooled cells were loaded onto a Chromium Next GEM Chip G with a total of 4 wells per pool, aiming to recover 5,000 cells per patient (2.6x$10^5$ total cells; except for pool #1 which had 2 wells per pool), and processed for single-cell encapsulation and cDNA library generation using the Chromium Single Cell 3′ v3.1 Reagent Kits (User guide CG000206 Rev D; 10X Genomics) and Single Index Kit T Set A (10X Genomics) according to the manufacturer's instructions. TotalSeq-A library generation was performed according to manufacturer's instructions (BioLegend). Gene Expression (GEX) and surface protein-targeting Antibody Derived Tag (ADT) libraries were pooled at 2.5:1 and the libraries were sequenced on Illumina NovaSeq 6000 (Illumina) with a target of 25,000 reads per cell for GEX and 10,000 reads per cell for ADT (S4-200, ~2.5B reads per library) at the Center for Advanced Technology at UCSF using the recommended cycle numbers: Read 1 28 cycles, i7 index 8 cycles, Read 2 91 cycles.

**CITE-seq data pre-processing, demultiplexing, doublet detection, and quality control**

Sequencer-obtained bcl files were demultiplexed using mkfastq program from Cell Ranger v6.0.2 (10x Genomics). Feature-barcode matrices were obtained for each library by aligning the raw fastqs of gene expression to GRCh38 reference genome (Ensembl v98) and the raw fastqs of surface protein expression to the TotalSeq$^{TM}$-A antibody reference panel (data file S3) using the Cell Ranger count. Raw feature-barcode matrices were imported and analyzed using Seurat (v4.0.3) (*62*).

Each library was demultiplexed to map individual cells to their sample of origin using freemuxlet (vAug2021) (*63*), SNP profiles were mapped to the genotypes of patients from bulk RNA-seq using bcftools gtcheck (v1.10.2)(*64*), and inter-sample doublets and empty droplets were removed. DoubletFinder (v2.0.3) (*65*) was used to filter barcodes containing heterotypic intra-sample doublets, as described (*66*), with default parameters except as specified: npcs=35, expected rate of intra-sample doublets = rate of inter-sample doublets (freemuxlet) divided by the number of samples in a pool. Barcodes containing less than 100 genes and genes that are detected in fewer than 3 cells were removed. We further filtered out the barcodes based on library-specific cutoffs for the number of genes per cell (nFeature_RNA <300-400 and >4,000), the number of unique molecular indexes (UMIs) per cell (nCount_RNA <700-2,000 and >20,000; nCount_ADT >20,000), mitochondrial content (>20%), and ribosomal content (<2-13% and >60%) to remove poor quality cells. Library SCG27 was removed due to abnormally high ribosomal content. The resulting high-quality cell-barcodes were used for further analysis.

An initial review of ADT reads revealed 28 out of 186 Total-seq-A exhibited high background/non-specific expression and were removed from clustering and UMAP analysis (data file S3). Preliminary clustering using a weighted nearest neighbor approach (described below) was used to identify and remove low-quality cells that lacked useful markers or were identified as red blood cells (defined as > 1 UMI count of HBB, HBA1, and/or HBA2 genes) or platelets (>1 UMI count of PPBP, PF4, CAVIN2, and/or GNG11 genes), comprising 6.5% of total cells. The resulting dataset of 519,228 cells was used for the following analyses.

**CITE-seq normalization, dimensionality reduction, and clustering**

Gene expression data from all libraries were merged, log-normalized, and scaled per cell with regression on UMI counts, mitochondrial content, ribosomal contents, and predicted cell-cycle state. The top 2,000 variable features were identified using the 'vst' method, PCA was performed using npcs=50, and batch-effect (library-specific bias) was normalized using Harmony (*67*).

ADT data was normalized and denoised using the 'denoised and scaled by background' (dsb) (*68*) method for each library separately using empty droplets as background, as defined by barcodes containing less than 100 UMI and more than 10 ADT counts. Since the antibody panel did not include isotype control antibodies, we did not perform isotype control antibody-based normalization. The normalized values smaller than -10 were truncated by substituting them with zero. We used reciprocal PCA to batch-correct and integrate the protein data. Briefly, after scaling the data and calculating principal components (PC) for each library, the data was integrated using IntegrateData function in Seurat. The integrated protein data was scaled, the ADT counts per cell and cell-cycle states were regressed out, and PCA was performed using npcs=30.

A weighted nearest neighbor (WNN) graph was constructed using the first 30 harmony dimensions for gene expression and first 18 integrated PCs for protein expression using Seurat's FindMultiModalNeighbors function with default parameters except prune.SNN = 1/20. Using the WNN graph, dimensionality reduction was performed using Uniform Manifold Approximation and Projection (UMAP) and clustering of single cells using smart local moving (SLM) algorithm in Seurat's FindClusters function. We generated clusters using resolution values between 0.5 and 4 and, based on manual inspection, selected the resolution of 2 to generate 51 clusters of cells. Separately, the raw surface protein expression data was transformed using centered log-ratio (CLR) for cluster annotation purposes.

#### CITE-seq cell annotation and differential expression

For cluster annotation, marker genes and proteins were identified for each cluster using the Model-based Analysis of Single Cell Transcriptomics (MAST) (*69*) algorithm with library as a latent variable using Seurat's FindMarkers and FindAllMarkers functions. We identified multiple clusters with mixed cell identities that did not separate into distinct clusters at any resolution (e.g., clusters with both CD4+ CD8- cells and CD4- CD8+ cells based on protein and gene counts), in part due to high background signal from non-specific antibodies (e.g., CD41, CD63, CD36), a common feature of ADT data. To separate these clusters of mixed identities into individual sub-populations, they were separately clustered using 0.2 or 0.3 resolution and the phenotypically distinct sub-populations were assigned new cluster identities. Based on the expression of canonical markers, we generated two sets of cluster annotations, one for broad cell types and another for finer cell subsets, resulting in 21 coarse populations representing individual immune cell types, including naïve and effector/memory T cells (Fig. 6A), and 34 clusters representing different subsets of immune cell types, respectively (fig. S8B and C). The 4.3% of cells lacking distinct gene expression patterns were left unlabeled. The reporting of differential expression analysis results was restricted only to coarse populations because the higher number of cells in coarse populations aided in reducing the noise introduced by sampling variability

#### Differential frequency analysis

The cell counts for each cell type were normalized by the total cells in the sample and multiplied by 100 to obtain percentages. The normalized cell frequencies were compared between groups of interest using Wilcoxon Rank-Sum test and the p-values were adjusted using Benjamini-Hochberg (BH) procedure with a cut-off of adj. $p<0.1$ deemed significant.

#### Differential expression analysis

To identify outcome-specific signal amidst a strong patient-specific effect revealed by hierarchical clustering, we leveraged the serial nature of this study to perform differential expression analysis using the limma package (v3.48.3) (*35*) and including paired samples from baseline and another timepoint-of-interest in the same analysis. Importantly, we used limma's duplicateCorrelation function to account for within patient correlation. We first summed the UMI counts across single cells for each sample, such that we obtained one sample x gene matrix per cell type. We subset the matrix to include samples from baseline and another timepoint of interest; we included only those patients that had both timepoints collected. We removed the lowly expressed genes by retaining only genes that had counts-per-million greater than 1 in more than 10% of samples. We included batches (pool ids), timepoint, and outcome variables in the model design, transformed the raw counts using limma's voom function, calculated patient-based correlation using duplicateCorrelation, and fitted linear models using lmFit function while using the patient-based correlation as input for 'correlation' parameter and patient id for 'block' parameter. We then computed log-odds of differential expression by empirical Bayes moderation of the standard errors towards a global value using eBayes function and considered genes with FDR < 10% as significant (data file S1). The following four samples were included in two independent pools of samples for 10X processing, XVIR1-HS41-SNEGB1, XVIR1-HS41-SNEGB6, XVIR1-HS42-SNEGB3, XVIR1-HS7-SNEGB4; for the paired differential gene expression analysis described above, the pool containing the most cells was used. We repeated this process for Effector/Memory CD4+ and Effector/Memory CD8+ T cells from iVR and Peak ALT.

Gene-set enrichment analysis (GSEA) (*39*) analysis was performed for Effector/Memory CD4+ and Effector/Memory CD8+ T cells from iVR, and Peak ALT, respectively, using KEGG, Hallmark, Reactome, GO biological process ontology gene set (C5) and Immunologic signature gene set (C7) from MSigDB (Broad Institute) databases. Briefly, we ranked the genes by the $\log_2$ fold-difference between clearance and persistence and evaluated the enrichment of each pathway using fgsea R package (v1.18.0). The pathways with adjusted p-value < 0.1 were considered significant (data file S2).

#### Statistical analysis

Unless otherwise specified, statistics were performed using Prism (Graph Pad Software). Statistical significance was determined by two-tailed unpaired Student's *t* test (when two groups were compared), two-tailed paired Student's *t* test (for paired data on the same mice), one-way ANOVA with Dunnett's correction for multiple comparisons to a control group (when one parameter was compared between more than two groups and a defined control group), or one-way ANOVA with Tukey's multiple-comparison test (when one parameter was compared for more than two biological groups, and all groups were compared to each other), or two-way ANOVA with Dunnett's correction for multiple comparisons to a control group (when multiple parameters were compared between more than two groups and a defined control group), or two-way ANOVA with Tukey's multiple-comparison test (when one parameter was compared for more than two biological groups, and all groups were compared to each other). For significance of HBsAg clearance, in which one of two possible outcomes was compared (clearance=1 versus no clearance=0), log-rank (Mantel-Cox) Chi-square test was used. For ordinal or ranked data (e.g., histology scores), the Mann-Whitney rank sum test was used. For CyTOF data multiple unpaired t-tests with Benjamini Hochberg FDR correction was used. In all figures with multiple *n*, data are presented as means ± SEM. $P < 0.05$ was considered significant. Statistical analysis for differential gene expression of CITE-seq data is described in the CITE-seq data analysis section.
